# Crucial Role of *Salmonella* Genomic Island 1 Master Activator in Parasitism of IncC plasmids

**DOI:** 10.1101/2020.08.27.269225

**Authors:** Romain Durand, Kévin T. Huguet, Nicolas Rivard, Nicolas Carraro, Sébastien Rodrigue, Vincent Burrus

## Abstract

IncC conjugative plasmids and the multiple variants of *Salmonella* Genomic Island 1 (SGI1) are two functionally interacting families of mobile genetic elements commonly associated with multidrug resistance in *Gammaproteobacteria*. SGI1 and its siblings are specifically mobilised *in trans* by IncC conjugative plasmids. Conjugative transfer of IncC plasmids is activated by the plasmid-encoded master activator AcaCD. SGI1 carries five AcaCD-responsive promoters that drive the expression of genes involved in its excision, replication, and mobilisation. SGI1 encodes an AcaCD homologue, the transcriptional activator complex SgaCD (also known as FlhDC_SGI1_) that seems to recognise and activate the same SGI1 promoters. Here, we investigated the relevance of SgaCD in SGI1’s lifecycle. Mating assays revealed the requirement for SgaCD and its IncC-encoded counterpart AcaCD in the mobilisation of SGI1. An integrative approach combining ChIP-exo, Cappable-seq, and RNA-seq confirmed that SgaCD activates each of the 18 AcaCD-responsive promoters driving the expression of the plasmid transfer functions. A comprehensive analysis of the activity of the complete set of AcaCD-responsive promoters in both SGI1 and IncC plasmid was performed through reporter assays. qPCR and flow cytometry assays revealed that SgaCD is essential for the excision and replication of SGI1, and the destabilisation of the helper IncC plasmid.

## INTRODUCTION

Multidrug resistant bacteria are increasingly recognised as an economic burden and a global threat to public health (1). Their emergence is being fuelled by diverse mobile genetic elements including genomic islands and conjugative plasmids (2) that often carry many antibiotic resistance genes. A better understanding of the mechanisms promoting the dissemination of mobile genetic elements is thus urgently needed. Mobilisable genomic islands (MGIs), also referred to as Integrated Mobilisable Elements (IMEs), have recently been getting renewed attention as they are increasingly recognised as important contributors to the propagation of multidrug resistance (3–7). MGIs usually carry diverse cargos conferring antibiotic or heavy metal resistance, bacteriocin synthesis or resistance to phage infection (7–9). *Salmonella* Genomic Island 1 (SGI1) is a 42.4-kb MGI that confers resistance to ampicillin, chloramphenicol, streptomycin, sulfonamides and tetracycline (ACSSuT) (10). SGI1 variants, which often bear a class 1 integron with diverse sets of antibiotic resistance gene cassettes, are found integrated at the 3’ end of *trmE* (also known as *mnmE* or *thdF*) in the chromosome of several species of *Gammaproteobacteriaceae*, including human pathogens such as *Salmonella enterica* serovars, *Proteus mirabilis, Morganella morganii, Providencia stuartii* or *Klebsiella pneumoniae* (11–15). SGI1 and its variants are specifically mobilised *in trans* by the IncC and the closely related IncA conjugative plasmids (16). IncC plasmids are large, broad-host-range, and globally distributed plasmids that contribute to the propagation of multidrug resistance genes. For instance, IncC plasmids are frequently associated with New Delhi metallo-β-lactamase genes (*bla*_NDM_) that confer resistance against most β-lactams including carbapenems (17), and have been a key driver of antibiotic resistance dissemination in seventh pandemic *Vibrio cholerae* O1 strains in Africa (18). Despite a lower epidemiological success compared to IncC plasmids, IncA plasmids have been found responsible for the spread of the carbapenemase gene *bla*_VIM-1_ in several species of *Enterobacterales* in hospitalised patients in Italy (19).

Conjugative transfer of IncC plasmids is controlled by two loci, the *acr1-acaDC-acr2* region, and the *acaB* gene located near *traNC. acaDC* encodes the heteromeric complex AcaCD distantly related to SetCD, the transcriptional activator of transfer genes of the integrative and conjugative elements (ICEs) of the SXT/R391 family (20, 21). AcaCD and SetCD are distantly related to FlhCD, the transcriptional activator of flagellar operons in *Escherichia coli* (20, 22, 23). Expression of *acaDC* is driven from the promoter *P_acr1_* that is repressed by Acr1 and Acr2 (20, 24). Recently, Hancock *et al.* demonstrated that *P_acr1_* is activated by AcaB, a novel transcriptional activator that exhibits structural similarity to bacterial transcription factors from the ribbon–helix–helix (RHH) superfamily (25). Interestingly, *acaB* has been shown to be part of the AcaCD regulon, wherein AcaCD and AcaB promote mutual expression, generating a positive feedback loop that activates conjugation. AcaCD turns on a total of 18 AcaCD-activatable promoters driving the expression of major operons involved in the formation of the mating pore and initiation of conjugative transfer, as well as genes of unknown function (20, 26). Furthermore, AcaCD also activates the expression of genes carried by distinct families of MGIs, including five operons in SGI1 (7, 26–28). The functions of five AcaCD-activatable genes of SGI1 have been characterised. *xis* encodes the recombination directionality factor that facilitates the excision of SGI1 from the chromosome catalysed by Int (29). *rep* encodes the replication initiator protein that initiates SGI1 replication at the origin of replication (*oriV*) (30). *traN_S_, traH_S_* and *traG_S_* encode three type IV secretion system (T4SS) subunits that replace their respective counterparts TraNC, TraHC and TraG_C_ in the mating pore encoded by the helper IncC plasmid (31).

MGIs are usually thought of as relatively passive mobile genetic elements that await the arrival of a helper self-transmissible element (a conjugative plasmid or an ICE) to escape their quiescent state and ride along with their helper element. To transfer to a new host, MGIs first need to excise from the host’s chromosome, and next to be translocated through the mating pore encoded by their helper element. For both processes, IncC-mobilised MGIs take advantage of the AcaCD regulon (7, 20, 27). Surprisingly, SGI1 has regularly challenged the assumption of passivity of MGIs (3). SGI1 has notably been shown to actively reshape the mating pore encoded by IncC conjugative plasmids to enhance its own propagation (31). SGI1 also actively replicates and carries a functional toxin-antitoxin system (*sgiAT*) that enhances its stability when an IncC plasmid is concomitantly present in the host (30, 32). In addition, SGI1 has been reported to destabilise helper IncA and IncC plasmids (33), a mechanism only recently attributed to the activation of the replicative state of excised SGI1 (30). Remarkably, SGI1-K, a member of the SGI1 family, is unable to destabilise IncC plasmids (33). SGI1-K lacks the 3’ half of *traN_S_ (S005*) and two upstream genes, *sgaD* (*S007*) and *sgaC (S006)*, that code for an AcaCD ortholog complex named SgaCD (also known as FlhDC_SGI1_) (28, 34, 35). SgaC and SgaD share 79 and 46% identity with AcaC and AcaD, respectively. Although SgaCD was reported to activate all five AcaCD-activatable promoters of SGI1 and to complement the *acaDC* deletion of the IncC plasmid R16a, its biological role in SGI1’s lifecycle remains unclear (28). Deletion of *sgaDC* was reported to have no impact on the mobilisation of SGI1-C by the helper IncC plasmid R55 (27). SGI1-C is a variant that differs from SGI1 by its smaller integron conferring resistance to spectinomycin, streptomycin and sulfonamides (36). Furthermore, deletion of *acaDC* of the helper IncC plasmid R16a was reported to completely abolish both self-transfer and SGI1-C mobilisation (27, 28).

Nevertheless, the conservation of *sgaDC* in most members of the SGI1 family suggests an important role in their lifecycle rather than simple redundancy for activation of AcaCD-activatable promoters. In this study, we investigated the relevance of SgaCD. First, we showed using mating assays that SgaCD is important for SGI1 mobilisation by IncC plasmids. Using a combination of ChIP-exo, Cappable-seq and RNA-seq approaches, we compared transcriptional activation by SgaCD and AcaCD of AcaCD-responsive promoters in SGI1 and in a model IncC plasmid. A systematic assessment of promoter activities through β-galactosidase reporter assays allowed us to measure the differential response of these promoters to SgaCD and AcaCD complexes. Finally, we unveiled the crucial role of SgaCD in the excision and replication of SGI1, ultimately leading to the destabilisation of IncC conjugative plasmids.

## MATERIAL AND METHODS

### Bacterial strains and media

Bacterial strains, plasmids and genomic islands used in this study are described in Table 1. Strains were routinely grown in lysogeny broth (LB) at 37°C in an orbital shaker/incubator and were preserved at −75°C in LB broth containing 20% (vol/vol) glycerol. Antibiotics were used at the following concentrations: ampicillin (Ap), 100 μg/ml; chloramphenicol (Cm), 20 μg/ml; kanamycin (Kn), 50 μg/ml or 10 μg/ml for single copy integrants of *pOPlacZ;* nalidixic acid (Nx), 40 μg/ml; spectinomycin (Sp), 50 μg/ml; tetracycline (Tc), 12 μg/ml; rifampicin (Rf), 50 μg/ml. To induce expression from pBAD30 and pAH56, LB medium was supplemented with 0.2% L-arabinose or 0.1 mM isopropyl β-D-1-thiogalactopyranoside (IPTG), respectively. Conjugation assays were performed as described previously (31).

**Table 1.**
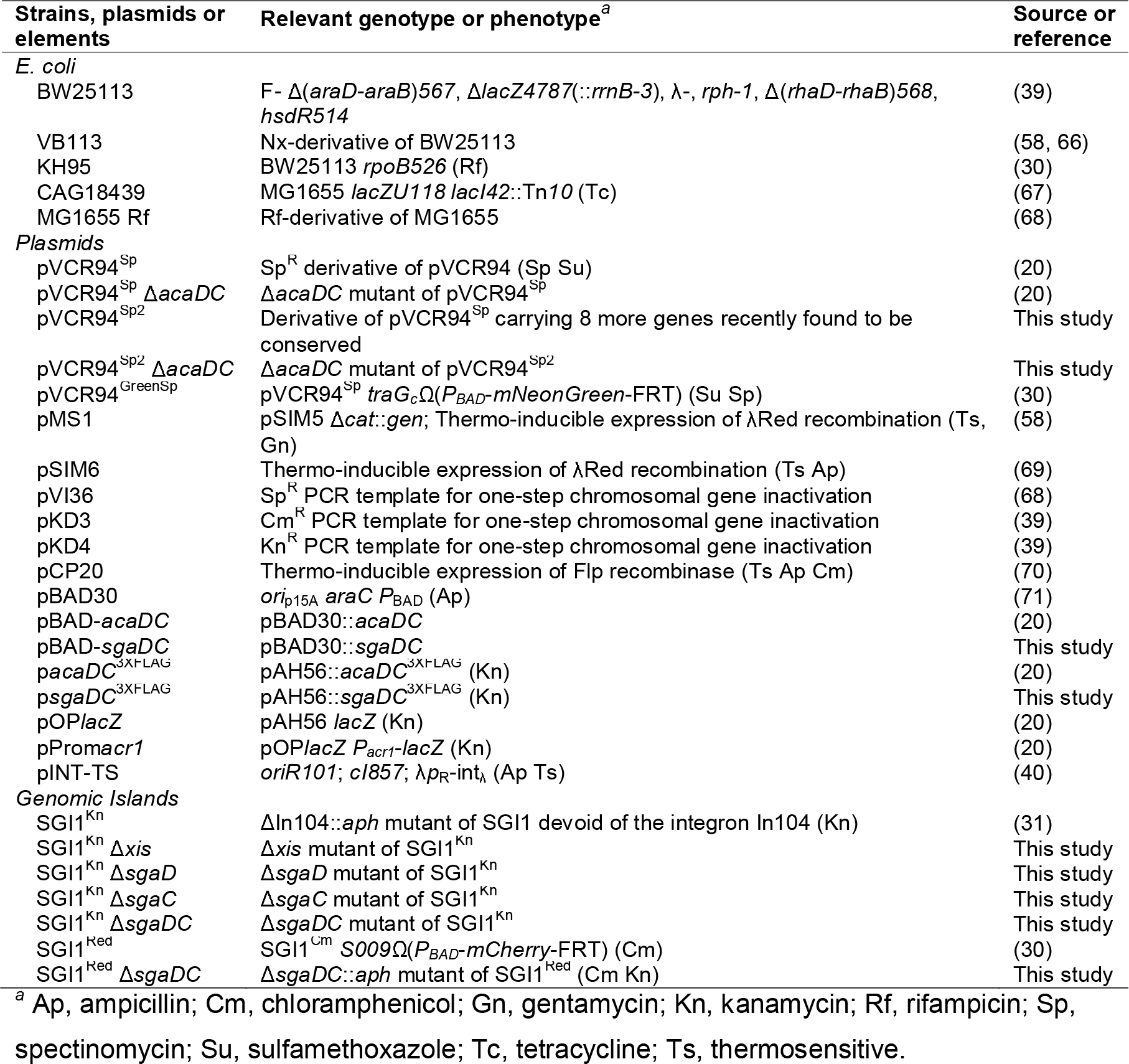
Strains and elements used in this study.

### Molecular biology

Plasmid DNA was prepared using either the EZ-10 Spin Column Plasmid DNA Miniprep Kit (Bio Basic) or the QIAprep Spin Miniprep Kit (Qiagen), according to manufacturer’s instructions. Genomic DNA was prepared using the QIamp DNA Mini Kit (Qiagen), according to manufacturer’s instructions. Restriction enzymes used in this study were purchased from New England Biolabs. Several DNA polymerases were used: Q5 (New England Biolabs), Taq (New England Biolabs) and Easy Taq (Civic Bioscience). PCR products were purified using either the EZ-10 Spin Column PCR Products Purification Kit (Bio Basic) or the QIAquick PCR Purification Kit (Qiagen), according to manufacturer’s instructions. *E. coli* was transformed by electroporation as described by Dower *et al.* (37) in a Bio-Rad GenePulser Xcell apparatus set at 25 μF, 200 Ω and 1.8 kV using 1-mm gap electroporation cuvettes. Sanger sequencing reactions were performed by the Plateforme de Séquençage et de Génotypage du Centre de Recherche du CHUL (Québec, QC, Canada).

### Plasmids and strains constructions

Oligonucleotides used in this study are listed in Supplementary Table S1. New sequencing data obtained from pVCR94 (NZ_CP033514.1) revealed that pVCR94^Sp^ (38) was missing eight conserved IncC genes. Therefore a new derivative, pVCR94^Sp2^, was constructed using the one-step chromosomal gene inactivation technique with pMS1, primer pair pVCR94delY2.f/94DelXnoFRT.rev and pVI36 as the template (39). Deletion mutants of pVCR94^Sp2^, SGI1^Kn^ and SGI1^Red^ were constructed in a similar fashion. Deletion of *acaDC* in pVCR94^Sp2^ was obtained using pSIM6, primer pair 94DelacaD.for/94DelacaC.rev and pKD4 as the template. Deletions of *xis, sgaD, sgaC* and *sgaDC* in SGI1^Kn^ were obtained using pSIM6, primer pairs SGI1delxis.for/SGI1delxis.rev, SGI1delsgaD07.for/SGI1delsgaD07.rev, SGI1delsgaC06.for/SGI1delsgaC06.rev and SGI1delsgaD07.for/SGI1delsgaC06.rev, respectively, and pKD3 as the template. Deletion of *sgaDC* in SGI1^Red^ was obtained using pMS1, primer pair SGI1delsgaD07.for/SGI1delsgaC06.rev and pKD4 as the template. When possible, the antibiotic resistance cassette was removed from the resulting construction by Flp-catalysed excision using pCP20. All deletions were verified by PCR and antibiotic resistance profiling. *sgaDC* was amplified using primer pair SGI1sgaDEcoRI.for/SGI1sgaCEcoRI.rev and genomic DNA of *E. coli* VB113 containing SGI^Kn^ as the template. The amplicon was then digested with EcoRI and cloned into EcoRI-digested pBAD30 using T4 DNA ligase (NEB), generating pBAD-*sgaDC*. p*sgaDC*^3XFLAG^ was derived from p*acaDC*^3XFLAG^. *sgaDC* was amplified using primer pair pAH56sgaCDinsF/pAH56sgaCDinsR and p*acaDC*^3XFLAG^ was linearized using primer pair pAH56sgaCDvecF/pAH56sgaCDvecR. These four primers were designed using NEBuilder® Assembly Tool (NEB). p*sgaDC*^3XFLAG^ was obtained by ligating both amplicons using the Gibson Assembly® Cloning Kit (NEB), to replace *acaDC* with *sgaDC.* PCR fragments containing the promoter region upstream of *vcrx012, vcrx035-36, vcrx059, vcrx068, traA, vcrx076, dsbC, traN_C_, acaB-vcrx087, vcrx098, vcrx114, vcrx128, vcrx140, S004* and *S018* were amplified using the corresponding primer pairs listed in Supplementary Table S1, and cloned into *pOPlacZ* using either PstI or PstI and XhoI to produce the corresponding *pOPlacZ* derivatives (Supplementary Figure S1). These vectors were ultimately integrated in single copy into the chromosomal site *attB_λ_* of *E. coli* BW25113 using pINT-Ts (40).

### ChlP-exo assays

LB medium supplemented with 50 μg/ml rifampicin, 10 μg/ml kanamycin, 50 μg/ml spectinomycin and, when necessary, 20 μg/ml chloramphenicol was inoculated with *E. coli* MG1655 Rf bearing p*sgaDC*^3XFLAG^, pVCR94^Sp2^ Δ*acaDC* and SGI1^Red^ Δ*sgaDC.* Induction of *sgaDC*^3XFLAG^ expression was done by adding 0.1 mM IPTG to cultures grown to an OD_600_ of 0.2, followed by a 1-h incubation at 37°C with shaking. 10 mL of culture was used for the ChIP-exo experiment, which was carried out as described previously (20), except for a shorter blocking step of magnetic beads used for immunoprecipitation: Dynabeads™ Protein A (Invitrogen™) were blocked by washing three times for 5 minutes each, in PBS/BSA (5 mg/mL). Replicates are detailed in Supplementary Table S2.

### Cappable-seq and RNA-seq assays

RNA was extracted from 5 mL cultures prepared as detailed above. Cultures were centrifuged for 10 min at 3,700 g, and the pellet was thoroughly resuspended in 1 mL TRI Reagent® (Sigma-Aldrich). Total RNA was then extracted using the Direct-zol RNA MiniPrep Kit (Zymo Research) with the recommended DNase I treatment, according to manufacturer’s instructions. Cappable-seq was carried out as described elsewhere (41), with the following modifications. Approximately 10 μg of clean RNA was used for each assay. The very first RNA clean-up step was performed using Agencourt® RNAClean® XP Beads (Beckman) to ensure maximum elimination of unincorporated DTB-GTP. The removal of 3’ phosphates from fragmented RNA was performed using the Thermo Scientific™ T4 Polynucleotide Kinase (Thermo Fisher Scientific) and its supplied ATP-free buffer. For RNA-seq samples, 800 ng of total RNA was fragmented in 5X RNA Fragmentation Buffer (200 mM Tris-Acetate, pH 8.1, 500 mM KOAc, 150 mM MgOA) by incubating at 95°C for 7 minutes and quenching immediately on ice. RNA was then purified using the RNA Clean & Concentrator-5 Kit (Zymo Research), according to manufacturer’s instructions. The quality and concentration of RNA before and after fragmentation were evaluated using a 2100 Bioanalyzer instrument (Agilent Technologies). Replicates are detailed in Supplementary Table S2.

### Illumina sequencing library preparation

ChIP-exo libraries were prepared as described previously (20), except for the second strand synthesis step that was performed using the Bst X DNA polymerase (Enzymatics) with ThermoPol® Buffer (NEB). Cappable-seq and RNA-seq libraries were prepared using the NEBNext® Small RNA Library Prep Set for Illumina® (NEB), according to manufacturer’s instructions, except NEBNext 5’ SR Adaptor for Illumina was replaced by previously described 5’-hybrid-A0 oligo (20). DNA molecules corresponding to the rRNA transcripts were depleted using the duplex-specific nuclease (Evrogen), as described elsewhere (42). All libraries were amplified and checked as previously described (20), then ultimately pooled. Illumina sequencing was performed on two different sequencing lines on a NextSeq® 500/550 High Output system at the Plateforme Rnomique de l’Université de Sherbrooke (Sherbrooke, QC, Canada).

### Bioinformatic analyses

Reads were trimmed with Trimmomatic (43) to discard nucleotides with a quality score below 30 and reads with a length below 36 bp. Quality was assessed before and after using FastQC (44). Trimmed reads were aligned on the *E. coli* MG1655 genome (NC_000913), pVCR94^Sp2^ Δ*acaDC* and SGI1^Red^ Δ*sgaDC* using Bowtie 2 (45). Alignment quality was assessed using SAMStat (46), and reads with a quality score below 10 were discarded using SAMtools view (47). Information on the number of reads before and after trimming, the quality of mapping and coverage for each sample, is detailed in Supplementary Table S2. Reads were then compressed, sorted and indexed using SAMtools (47). ChIP-exo and Cappable-seq reads were chopped to their first nucleotide and density was calculated separately for each DNA strand using BEDTools genomecov (48). Density files were ultimately compressed to BigWig format and visualised on the UCSC Genome Browser.

The footprint profile for each transcription start site of interest was analysed using the Versatile Aggregate Profiler (VAP) (49). Three pairs of divergent AcaCD-dependent promoters (*vcrx035-036, vcrx059-traI* and *acaB-vcrx087*) were excluded from the analysis to prevent potentially overlapping ChIP-exo signals on the positive and negative strands. Replicates of a given condition were pooled using SAMtools and reads were treated as described above. Densities for any condition and strand were normalised by the total signal obtained for a given reference (pVCR94^Sp2^ Δ*acaDC* or SGI1^Red^ Δ*sgaDC*). The start and end coordinates of each transcription start site of interest were used as reference points. The signal was reported for each base pair and represented as median and either 1^st^-9^th^ deciles or full range without smoothing. RPKM values were calculated for each DNA strand separately, using a script adapted from EDGE-pro (50). Differential expression analysis was performed using DESeq2 (51). Signal was also calculated for each condition on 1-kb intervals using BEDTools multicov (48). Log-transformed values were used to represent the variability between replicates (Supplementary Figure S2).

Search for SgaC homologues was carried out using Blastp (52) against the Genbank non-redundant protein sequence (nr) database restricted to the *Enterobacteriaceae* (taxid:543).

### β-galactosidase assays

The assays were carried out as described previously, using *o*-nitrophenyl-β-D-galactopyranoside (ONPG) as substrate (53). Cultures were prepared with LB medium supplemented with 10 μg/ml kanamycin to select the strain and 50 μg/ml ampicillin to maintain pBAD-*acaDC* or pBAD-*sgaDC*. Induction of *acaDC* or *sgaDC* was done by adding 0.2% arabinose to a refreshed culture grown to an OD_600_ of 0.2, followed by a 2-h incubation at 37°C with shaking prior to cell sampling.

### qPCR assays

Genomic DNA was obtained from 1 mL of cell cultures of *E. coli* VB113 bearing pVCR94^Sp^, SGI1^Kn^ or their mutants, grown for 16 h in LB medium supplemented with 40 μg/ml nalidixic acid, 50 μg/ml spectinomycin and 50 μg/ml kanamycin. Genomic DNA purity and concentration were measured with an ND-1000 NanoDrop spectrophotometer (Thermo Fisher Scientific). *attB* (236 bp), *S026* (234 bp) from SGI1^Kn^, *repA* (237 bp) from pVCR94^Sp^, as well as reference genes *dnaB* (235 bp), *hicB* (235 bp) and *trmE* (238 bp), were quantified using primer pairs qAttBFw/qAttBRv, qS026Fw/qS026Rv, qFwpVCR/qRvpVCR, qdnaBFw/qdnaBRv, qhicBFw/qhicBRv and qthdFFw/qthdFRv, respectively (Supplementary Table S1). Excision of SGI1^Kn^ was calculated as the ratio of *attB* per chromosome, SGI1^Kn^ and pVCR94^Sp^ copy numbers were calculated as the ratios of *S026* and *repA* per chromosome, respectively. Normalisation was done using all three reference genes (54). For clarity, the strain VB113 bearing both pVCR94^Sp^ and SGI1^Kn^ was considered to exhibit a 100% excision rate. VB113 bearing either pVCR94^Sp^ or SGI1^Kn^ alone, were presumed to contain only one copy of a given element. qPCR experiments were performed in biological triplicate at the Plateforme Rnomique de l’Université de Sherbrooke (Sherbrooke, QC, Canada).

### Cohabitation assays

The assays were carried out as described previously (30). Culture samples were diluted 1:1,000 in 1 mL of PBS. Fluorescence intensity of NeonGreen and mCherry in cells was monitored by flow cytometry analysis on a BD FACSJazz (BD Biosciences), and data were acquired with the BD FACS Sortware. mNeonGreen and mCherry were excited with 488 and 561 nm solid-state lasers, and their emission was detected using 513/17 and 610/20 nm emission filters, respectively. For each sample, fluorescence of 20,000 cells was captured, and the data was analysed using FCS Express 7 (De Novo Software).

### Statistical analyses and figures

Prism 8 (GraphPad Software) was used to plot graphics and to carry out statistical analyses. All figures were prepared using Inkscape 1.0 (https://inkscape.org/).

## RESULTS

### SGI1 rescues the transfer of an IncC plasmid lacking its master activator of transfer

Given the similarity between AcaCD and SgaCD, and the essentiality of AcaCD for IncC plasmid transfer activation, we first assessed the importance of *sgaDC.* Transfer rates of SGI1^Red^ and the coresident IncC helper plasmid pVCR94^Sp2^ were assessed in mating assays using combinations of wildtype and Δ*DC* mutants of both elements. Deletion of *acaDC* had no impact on transfer of the helper plasmid, SGI1^Red^ or cotransfer of both elements, confirming that SGI1 can complement the loss of *acaDC* (Figure 1A). In contrast, deletion of *sgaDC* reduced transfer of SGI1^Red^ nearly 2,000-fold compared to the wildtype, whereas it had no impact on transfer of the helper plasmid or cotransfer of both elements, showing that *sgaDC* is important for SGI1 transfer and/or stability (Figure 1A). Deletion of both *acaDC* and *sgaDC* abolished transfer of both elements. Finally, overexpression of a single chromosomal copy of *sgaDC*^3XFLAG^ under the control of the IPTG-inducible *P_tac_* promoter activated transfer of both elements beyond wildtype levels (Figure 1A). To confirm that this phenotype was exclusively attributable to SgaCD, *trans*-complementation with chromosomal *P_tac_-sgaDC*^3XFLAG^ was also performed using donors lacking SGI1. While deletion of *acaDC* abolished pVCR94^Sp2^ transfer in this context, it was fully restored by *sgaDC*^3XFLAG^ overexpression (Figure 1B).

**Figure 1.**
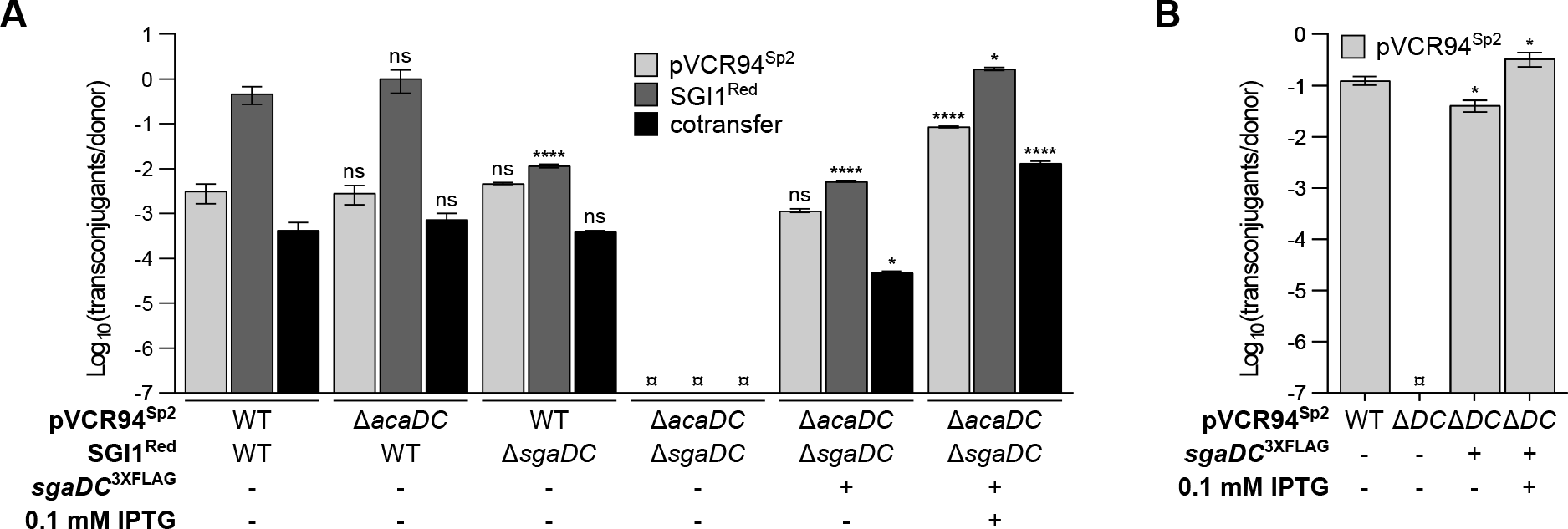
Role of SgaCD in conjugative transfer of IncC plasmids and SGI1. Effect of *sgaDC* on conjugative transfer of pVCR94^Sp2^ (light grey bars) in the presence (A) or absence (B) of SGI1^Red^ (dark grey bars). Conjugation assays were carried out using MG1655 Rf containing the indicated elements as donor strains and *E. coli* CAG18439 (Tc) as the recipient strain. Wild-type (WT) or derivative mutants of both elements are indicated under each graph. For clarity, *acaDC* is shortened *DC* in (B). Transfer frequencies are expressed as the number of transconjugants per Rf^R^ Sp^R^ Cm^R^ donor CFUs in (A), or Rf^R^ Sp^R^ donor CFUs in (B). The bars represent the mean and standard error of the mean obtained from a biological triplicate. ¤ indicates that the frequency of transfer was below the detection limit (<10^−7^). For each panel and each element, a one-way ANOVA with Dunnett’s multiple comparison test were performed on the logarithm of the values to compare each bar to the WT control (first bar). Statistical significance is indicated as follows: ****, *P* < 0.0001; ***, *P* < 0.001; **, *P* < 0.01; *, *P* < 0.05; ns, not significant.

### SgaCD activates all AcaCD-activatable promoters and generates the same binding footprint as AcaCD

The SgaCD regulon was characterised using a combination of ChIP-exo, Cappable-seq and RNA-seq approaches to identify the binding sites, transcriptional start sites (TSS) and quantify mRNA transcript levels, respectively. In these experiments, either *sgaDC*^3XFLAG^ or *acaDC*^3XFLAG^ was expressed from *P_tac_* instead of the native promoter to ensure homogenous expression in cell populations. Otherwise isogenic cells lacking *sgaDC*^3XFLAG^ and *acaDC*^3XFLAG^ were used as negative controls. We also used Δ*DC* mutants of pVCR94^Sp2^ and SGI1^Red^ to prevent interference from the native unlabelled complexes. The ChIP-exo experiment revealed that SgaCD binds all known AcaCD-activatable promoters on pVCR94^Sp2^ and SGI1 (Figure 2). No additional promoter was found to be bound by SgaCD, showing that AcaCD and SgaCD complexes have similar activities and recognise identical sequence motifs. ChIP-exo peaks usually paired with those of Cappable-seq on the positive DNA strand, except for the control condition, confirming that SgaCD binding correlates with transcription of each promoter (Figure 2). Transcriptomic data obtained from RNA-seq assays with pVCR94^Sp2^ confirmed that *sgaDC* expression leads to an overall increase of mRNA levels, especially for operons associated with conjugative transfer functions (Figure 2). Likewise, expression of most SGI1 genes increased upon expression of *sgaDC* or *acaDC* (Figure 2).

**Figure 2.**
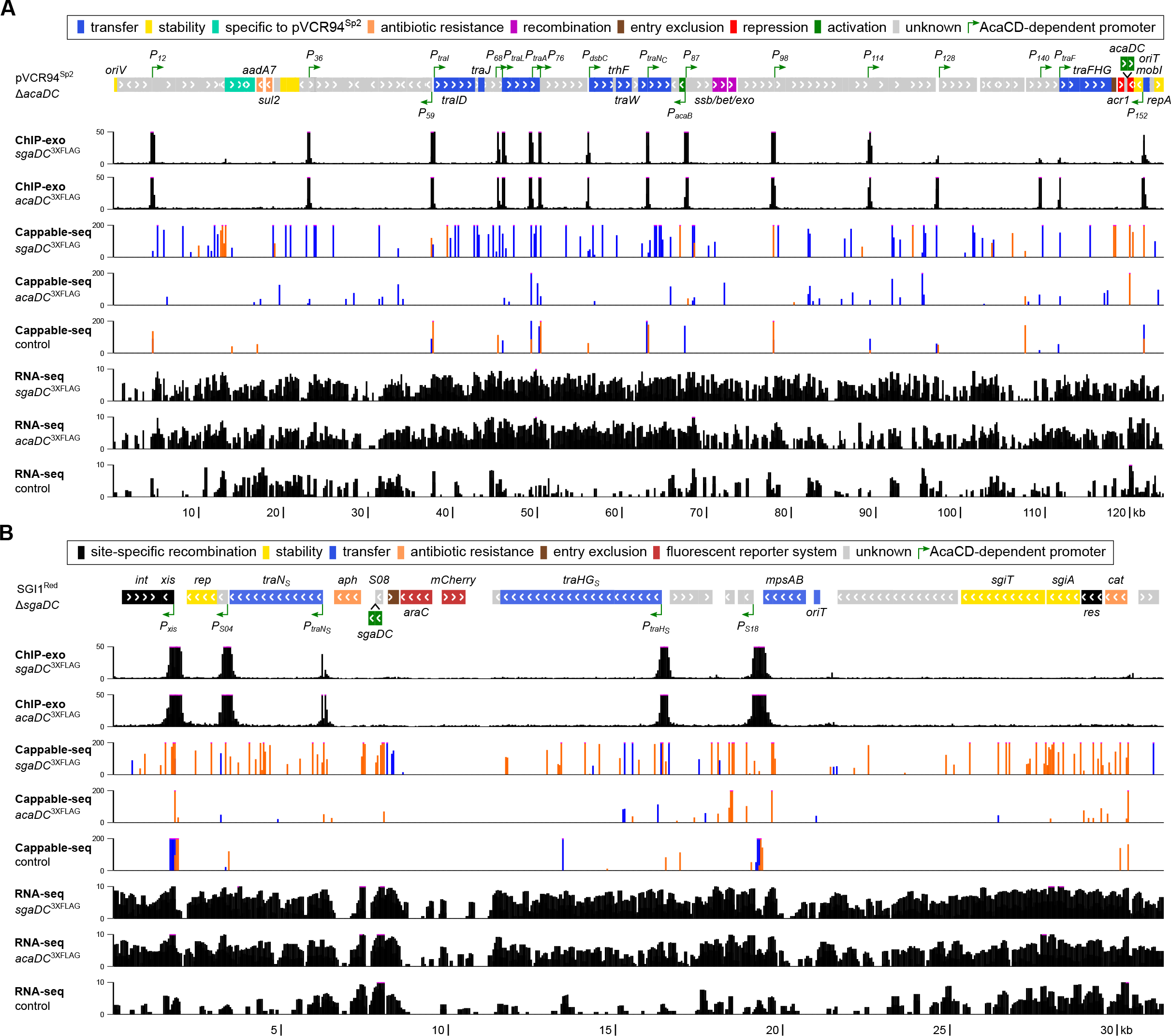
In-depth analysis of the AcaCD/SgaCD regulon. Results of ChIP-exo, Cappable-seq and RNA-seq experiments on *E. coli* MG1655 Rf carrying both pVCR94^Sp2^ Δ*acaDC* (A) and SGI1^Red^ Δ*sgaDC* (B) with or without a single chromosomal copy of either p*sgaDC*^3XFLAG^ expressing the native SgaD subunit along with a C-terminal 3XFLAG-tagged SgaC subunit induced by IPTG, or p*acaDC*^3XFLAG^. The circular map of the plasmid was linearised after the *repA* gene. The location and orientation of ORFs are depicted by arrowed boxes, which are color-coded by function as indicated. Green arrows indicate previously identified (A) or predicted (B) AcaCD-dependent promoters. Each track (one representative replicate per condition as detailed in Supplementary Table S2) plots the number of mapped reads as a function of the position in each element. Read densities are displayed as black bars, or blue and orange bars for Cappable-seq densities on the positive and negative DNA strands (linear scale for ChIP-exo and Cappable-seq densities, log scale for RNA-seq densities). Pink dots at the summit of peaks indicate a signal beyond the represented y-axis maximal value.

To compare the binding footprint of both complexes, two aggregated profiles were built by compiling ChIP-exo and Cappable-seq signals of pVCR94^Sp2^ and SGI1 promoters. No major difference could be observed between the SgaCD and AcaCD profiles (Figure 3 upper panels). Both complexes bind to the same site located −64 to −37 bp upstream of the transcription start site (TSS) (20). The large protected sequence starts −42 bp upstream of the TSS, that is 4 bp downstream of the GCCCNDWWWGGGC palindromic motif and ends 22 bp after the TSS. This protection is compatible with the promoter-bound RNA polymerase holoenzyme complex, as previously reported for class II activation where the bound activator complex abuts the promoter −35 element (Figure 3 upper panels and Supplementary Figure S3A-E) (20, 21, 55, 56). The sequence overlapping the binding motif, where two peaks are observed immediately after `GCCC` and `GGGC`, corresponds to the AcaCD/SgaCD footprint. The AcaCD profile displayed sharper boundaries than that of SgaCD with a stronger signal at these positions (Figure 3 upper panels and Supplementary Figure S3A-E). Assuming comparable crosslink efficiencies, this observation suggests better recruitment and tighter binding of AcaCD compared to SgaCD. Remarkably, the Cappable-seq signal of SgaCD on SGI1 promoters was weaker than that of AcaCD. Nevertheless, these results show that SgaCD and AcaCD, though encoded by two unrelated mobile genetic elements, recognise and bind identical sequence motifs.

**Figure 3.**
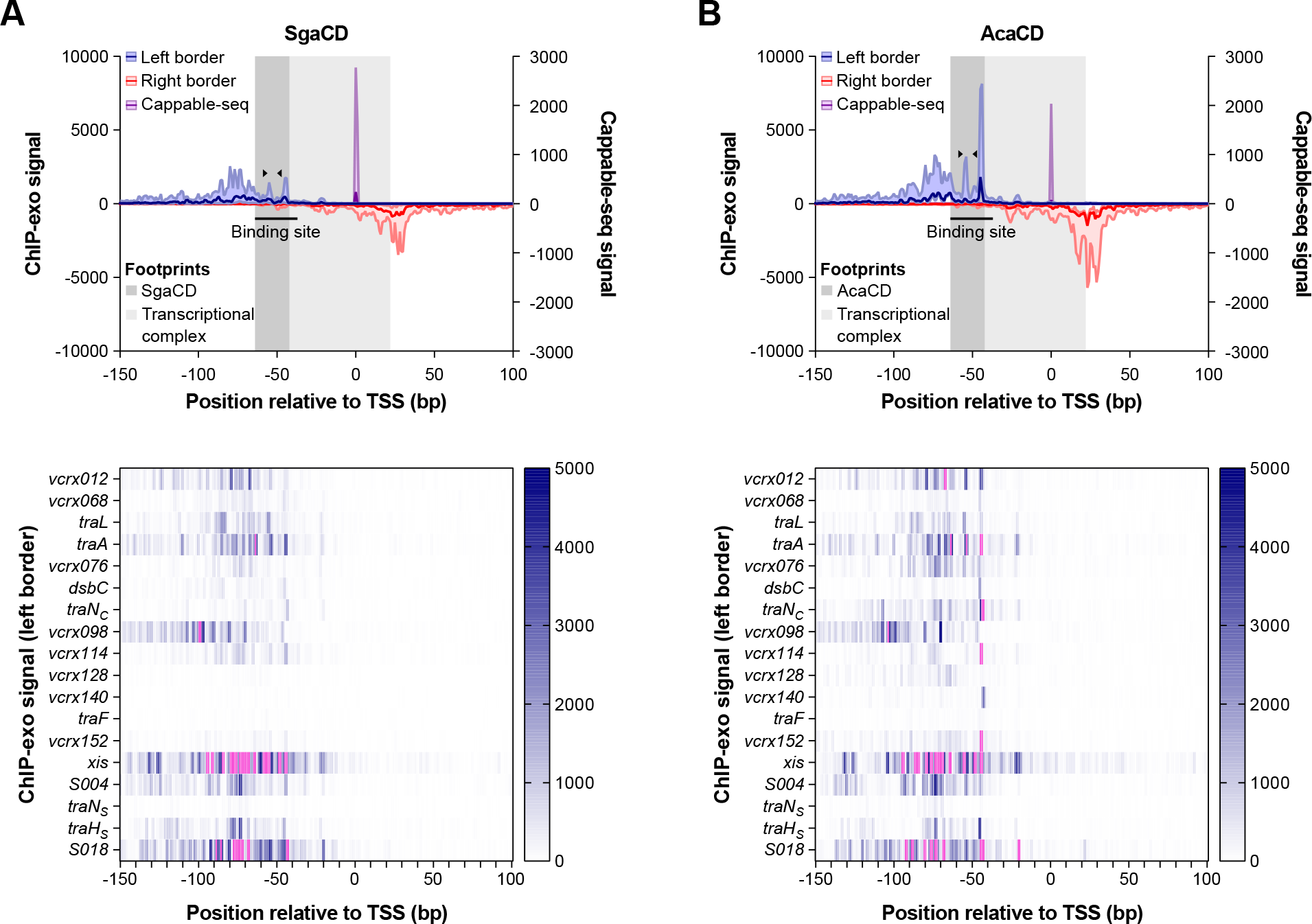
VAP profiles of AcaCD/SgaCD footprint. Upper panels: ChIP-exo and Cappable-seq profiles obtained with SgaCD (A) and AcaCD (B) were aggregated from 13 AcaCD-activatable IncC promoters and all 5 AcaCD-activatable SGI1 promoters. The normalised density of ChIP-exo reads mapping on the positive DNA strand (left border, in blue) and the negative strand (right border, in red), as well as the normalised density of Cappable-seq reads mapping on the positive strand (purple), are plotted as functions of the distance in nucleotides to the aggregated transcription start site (TSS). Data are represented as median, first and ninth deciles. The binding site is indicated as a black line, with its **GCCC**NDWWW**GGGC** motif delimitated by black triangles. The region protected by the transcriptional complex, delimitated by the last and first peaks on the ChIP-exo left and right borders, respectively, is depicted in light grey. The region protected by either SgaCD or AcaCD, starting at the binding site, is depicted in dark grey. Lower panels show the ChIP-exo signal (left border) for each promoter as a heatmap, where pink indicates a signal beyond 5,000.

Both complexes appeared to bind better to *P_xis_* relatively to other AcaCD-activatable promoters (Figure 3 lower panels). Comparison of individual profiles obtained for *P_xis_* and *P_S004_* promoters with AcaCD and SgaCD, revealed clear differences. AcaCD and SgaCD profiles at *P_xis_* were similar to the aggregated profiles (Supplementary Figure S3E-F), supporting the hypothesis that SgaCD binds less efficiently than AcaCD, leading to a weaker transcription initiation, as suggested by the weaker Cappable-seq signal. In strong contrast, the individual profiles obtained with SGI1-borne *P_S004_* promoter were unique (Supplementary Figure S3G-H). Comparison of ChIP-exo signals revealed efficient binding of SgaCD but not AcaCD. Detection of a strong AcaCD ChIP-exo signal on the right border shortly after the binding site suggests premature detachment or poor recruitment of the transcriptional activator (Supplementary Figure S3H). The extremely weak Cappable-seq signal supports the hypothesis of a faulty transcriptional initiation by AcaCD. Except for a stretch of five Gs located between the binding site and −10 element, no specific feature could be found that could explain the weak activation of *P_S004_* by AcaCD (Supplementary Figure S1). In contrast, the Cappable-seq signal obtained with SgaCD was higher and displayed 3 peaks, at the predicted, and 5 and 8 bp after the TSS. In summary, SgaCD seems to activate *P_S004_* with much higher efficiency than its IncC-encoded homolog.

### Overexpression of *sgaDC* significantly affects the transcriptome

A differential expression analysis was performed on transcriptomic data to identify up- and down-regulated genes following *sgaDC* overexpression. This analysis considers the different sequencing depths of each condition and provides statistical metrics. A vast majority of differentially expressed genes were found to be activated by SgaCD (Figure 4A). For instance, in the helper IncC plasmid, the expression of *traL* or *traK* that encode predicted inner and outer membrane subunits of the conjugal T4SS, respectively, increased 500 to 1,000-fold (Figure 4A). In contrast, few genes were found to be downregulated upon *sgaDC* overexpression. The most significant reduction of expression was observed for *vcrx119* and *vcrx120*, two genes of unknown function. Since no binding signal was detected upstream of these two genes, repression by SgaCD is likely indirect (Figure 2). Furthermore, expression of the mobilisation gene *mobI*, which encodes a key factor for initiation of conjugative transfer at *oriT*, remained unchanged, confirming its independence from AcaCD and SgaCD (Figure 4A) (20, 57, 58). Finally, none of the 8 genes of pVCR94^Sp2^ that are absent in pVCR94^Sp^ were expressed under the control of an AcaCD-activatable promoter (Figures 2 and 4A). 916 chromosomal genes were differentially expressed upon *sgaDC* overexpression (Supplementary Figure S4 and Supplementary Table S3). Several of the most up-regulated genes have been shown to be involved in resistance to stress and membrane transport. Down-regulated genes are involved in cysteine and enterobactin synthesis, outer membrane transport and anaerobic respiration. As no SgaCD binding could be detected by ChIP-exo in the promoter region of the affected genes, SgaCD influence on chromosomal gene expression was likely indirect. Four additional differential expression analyses were conducted to evaluate the impact of SGI1 and to compare activation by SgaCD and AcaCD. Expression of *mobI* was repressed upon overexpression of *sgaDC* in the presence of SGI1^Red^ Δ*sgaDC* (Figure 4B). Since expression of *mobI* is independent of SgaCD, such repression likely results from expression of an SGI1-encoded factor. Few genes were found to be differentially expressed when overexpressing *sgaDC* compared to *acaDC* (Figures 4C and 4E). Therefore, a weaker transcriptional initiation as suggested by Cappable-seq does not necessarily lead to lower levels of transcripts in the artificial context of overexpressing these transcriptional activators. Finally, we confirmed that most SGI1 genes are up-regulated by SgaCD, *S004* and *traH_S_* presenting the highest fold-change values (Figure 4D).

**Figure 4.**
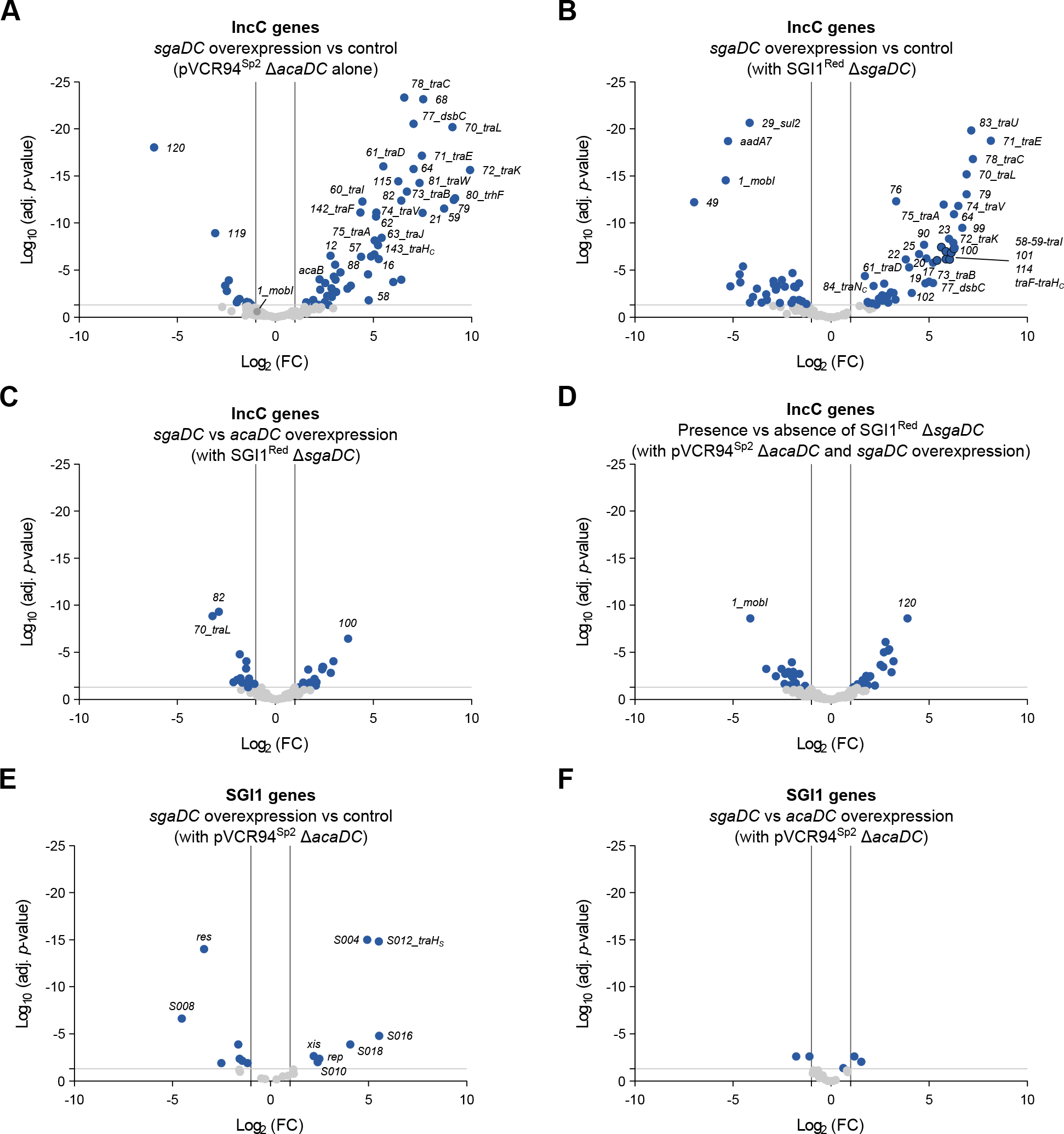
Differential expression analysis. Volcano plot of IncC (A-D) and SGI1 (E-F) genes using the Log_2_ of fold-change and adjusted *p*-value. Conditions are described at the top of each panel. The vertical lines indicate a 2-fold change in expression, and the horizontal line demarcates statistical significance at *p* = 0.05. Differentially expressed genes are marked as blue dots. For clarity, IncC genes are labeled “*001*” instead of *“vcrx001”.*

### Activation of gene expression by SgaCD is comparable to AcaCD

Since transcriptomic data only show the outcome of potentially multiple regulatory processes, ChIP-exo and Cappable-seq results were confirmed using an expression reporter assay based on transcriptional *lacZ* fusions to each of the 23 AcaCD-activatable promoters inserted in single copy into the chromosome. β-galactosidase assays were carried out with both AcaCD and SgaCD to quantify the relative expression level with each activator complex (Figure 5 and Supplementary Figure S1). While *vcrx036* and *vcrx087* were constitutively expressed, virtually all other promoters were directly activated by AcaCD and SgaCD (Figure 5A-C). Induction ratios of SgaCD were almost invariably lower than those of AcaCD, confirming previously reported results obtained with the 5 AcaCD-dependent promoters of SGI1 (Figure 5D-E) (28). However, the difference was not statistically significant for 15 out of 23 promoters. The gene of unknown function *vcrx087* was expressed both constitutively and under the control of AcaCD/SgaCD, which can be explained by the presence of nearly canonical σ^70^ −35 and −10 elements in its promoter sequence (Figure 5B and Supplementary Figure S1).

**Figure 5.**
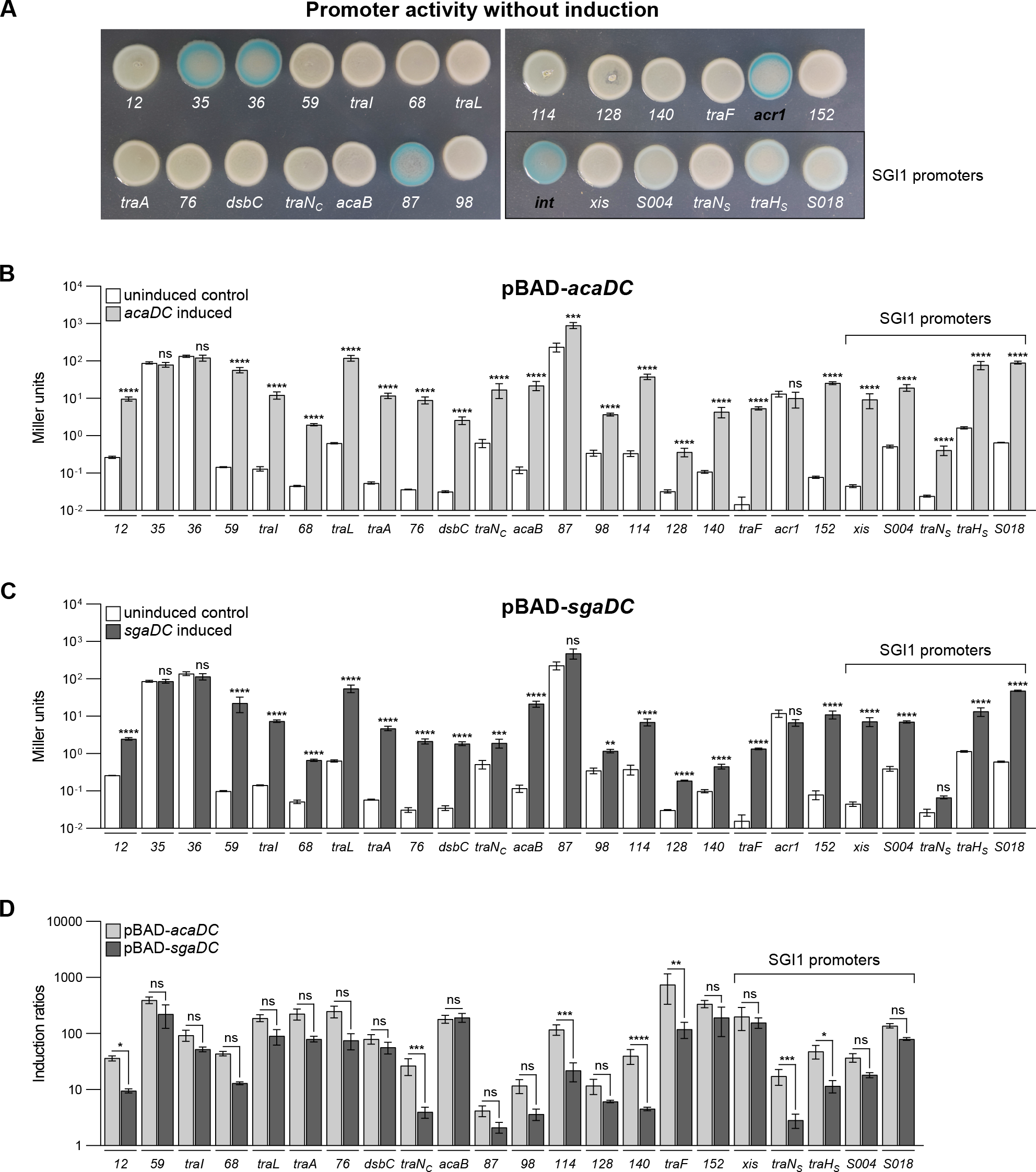
Systematic analysis of AcaCD/SgaCD-dependent promoters. Activity of all AcaCD-dependent promoters on IncC plasmids and SGI1 was monitored from single-copy, chromosomally integrated transcriptional *lacZ* fusions in *E. coli* BW25113. (A) Qualitative assay on LB medium supplemented with X-Gal. Promoters are identified by the first gene of the corresponding operon. AcaCD-independent promoters are labelled in black. (B, C) Miller units. (D) Induction ratios. β-galactosidase assays were carried out in LB medium supplemented with (grey bars in A, B) or without (white bars in A, B) arabinose to express *acaDC* or *sgaDC* from *P_BAD_* on pBAD-*acaDC* (A) or pBAD-*sgaDC* (B). The bars represent the mean and standard error of the mean obtained from a biological triplicate. In panel D, the x axis crosses the y axis at y = 1, indicating no induction. For panels B-D, three independent two-way ANOVA with Sidak’s multiple comparison test were performed on the logarithm of the values to compare the pair of bars for each promoter. Statistical significance is indicated as follows: ****, *P* < 0.0001; ***, *P* < 0.001; **, *P* < 0.01; *, *P* < 0.05; ns, not significant.

### SgaCD is essential for SGI1 replication

To investigate the role of SgaCD in SGI1’s lifecycle, we conducted qPCR assays using different mutants of pVCR94^Sp^ and SGI1^Kn^. As shown previously (20, 29), SGI1 excision was undetectable in the absence of the helper plasmid or when a Δ*xis* mutant was used (Figure 6A). Surprisingly, deletion of *acaDC* had a rather limited, statistically non-significant impact on the excision rate of SGI1^Kn^. On the contrary, deletion of *sgaDC* reduced the excision rate nearly 2,900-fold compared to the wild-type level, supporting a considerable role of SgaCD compared to AcaCD in SGI1’s lifecycle. Deletion of both resulted in the abolition of excision, indicating that residual excision of SGI1^Kn^ Δ*sgaDC* is triggered by the helper plasmid-encoded AcaCD. A comparable phenotype was observed when measuring the copy number of SGI1^Kn^ (Figure 6B). Deletion of *sgaDC* completely abolished SGI1 replication, whereas deletion of *acaDC* reduced it only 2.7-fold (Figure 6B). Together, these results account for the reduced mobilisation of SGI1^Kn^ Δ*sgaDC* by pVCR94^Sp^ (Figure 1A). None of the deletions had any statistically significant impact on pVCR94^Sp^ copy number (Figure 6C).

**Figure 6.**
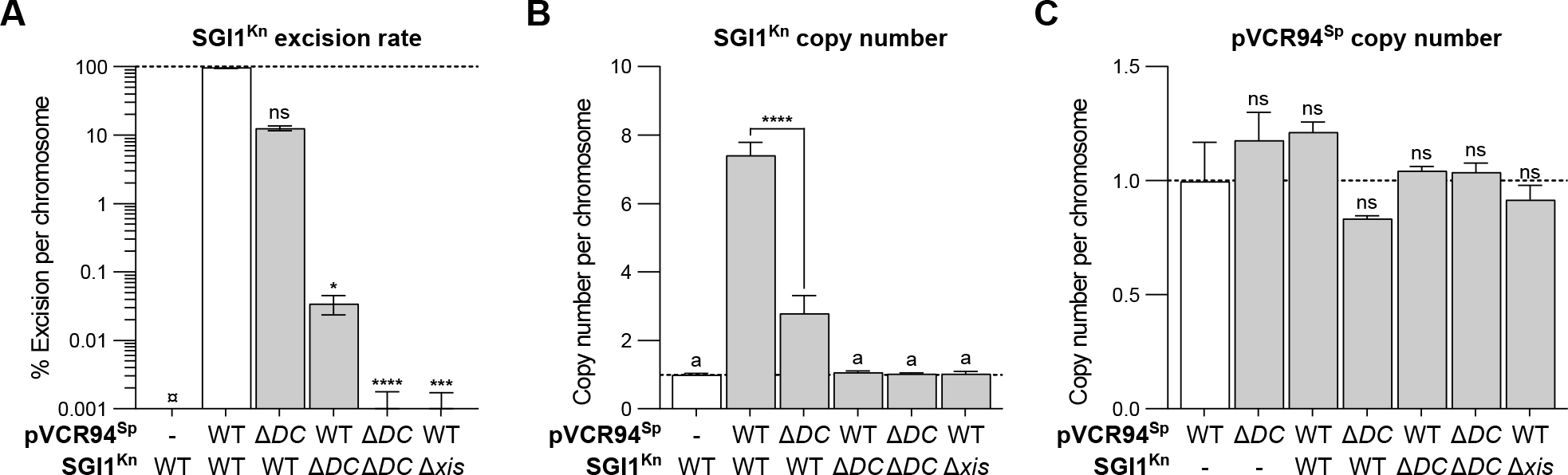
Effect of *sgaDC* and *acaDC* deletions on SGI1^Kn^ dynamics and pVCR94^Sp^ stability. All targets were amplified by qPCR alongside three reference genes: *trmE, hicB* and *dnaB.* (A) SGI1^Kn^ excision rate corresponds to the *attB*/chromosome ratio. (B) SGI1^Kn^ copy number corresponds to the *s026*/chromosome ratio. (C) pVCR94^Sp^ copy number corresponds to the *repA*/chromosome ratio. For each panel, all ratios were normalised using the control set to 1 and displayed in white. The bars represent the mean and standard error of the mean obtained from a biological triplicate. ¤ indicates that the excision rate was below the detection limit (< 0.001%). Statistical analyses were performed (on the logarithm of the values for panel A) using three independent one-way ANOVA with Tukey’s multiple comparison test. “a” indicates the bars showed no significant difference between one another. For panels A and C, statistical significance indicates comparisons to the normalisation control. Statistical significance is indicated as follows: ****, *P* < 0.0001; ***, *P* < 0.001; *, *P* < 0.05; ns, not significant.

### SgaCD-activated replication of SGI1 destabilises the helper IncC plasmid

Incompatibility between SGI1 and a coresident helper IncC plasmid has been shown to be linked to the replicative state of SGI1 (30). To test whether *sgaDC* had any role to play in SGI1 replication and incompatibility, we used a previously designed flow cytometry assay aimed at assessing the evolution of a population of cells carrying red fluorescence-producing SGI1^Red^ and green fluorescence-producing pVCR94^GreenSp^ over time. Cells producing red fluorescence segregate into two populations. Low-intensity red-fluorescent cells carry a single copy of SGI1 usually integrated in the chromosome, whereas cells producing high-intensity red fluorescence contain excised replicating SGI1 (30). pVCR94^GreenSp^ alone remained relatively stable and was retained in 71% of cells at T48 (Figure 7A). Cells carrying SGI1^Red^ or its Δ*sgaDC* mutant produced mostly low-intensity red fluorescence and remained steady (Figure 7B-C). Overexpression of *acaDC* was sufficient to promote SGI1 excision and replication in the absence of the helper IncC plasmid as 71% of the cells produced high-intensity red fluorescence at T0 (Figure 7D). However, SGI1^Red^ Δ*sgaDC* was rapidly lost as most cells failed to produce any fluorescence at T24. Overexpression of *sgaDC* resulted in a more widespread activation of SG1 replication as more than 90% of cells were highly red fluorescent at T0 (Figure 7E). SGI1 loss was delayed with *sgaDC* compared to *acaDC*, with more than 6% of cells retaining replicating SGI1 at T48. Together, these results show that SGI1 excision and replication can occur in the absence of the helper IncC plasmid when provided with AcaCD or SgaCD, although the latter seemed to promote stronger activation of *rep* expression. However, after excision, replication of SGI1 was insufficient to ensure its inheritance in the cell population in the absence of the IncC plasmid, despite the presence of the *sgiAT* toxin-antitoxin system.

**Figure 7.**
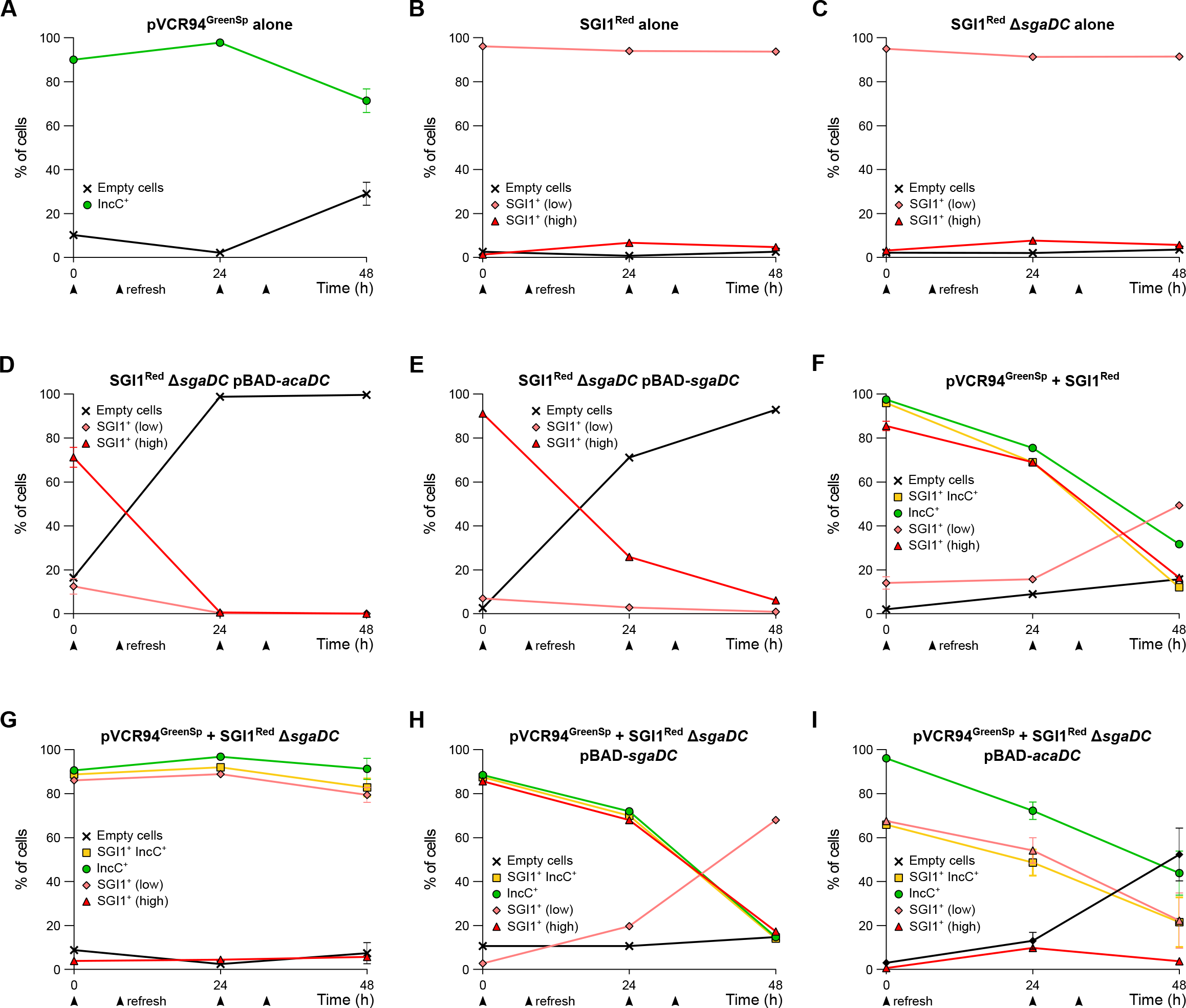
Effect of *sgaDC* deletion on incompatibility between SGI1 and IncC plasmids in the absence of selective pressure. Evolution of the percentage of *E. coli* KH95 cells bearing (A) pVCR94^GreenSp^ (IncC) or (B) SGI1^Red^ or (C-E) SGI1^Red^ Δ*sgaDC* (SGI1) or the indicated combination of both elements (F-I), in the absence or presence of pBAD-*sgaDC* (D, H) or pBAD-*acaDC* (E, I), over 48 hours in the absence of antibiotics (except for ampicillin for maintenance of pBAD derivatives) as monitored by flow cytometry. Plots show the mean and standard error of the mean values obtained from a biological triplicate.

When SGI1^Red^ was in the presence of pVCR94^GreenSp^, most cells produced high red fluorescence at T0, confirming SGI1 replication (Figure 7F). At T48, only 16% of the cells produced high red fluorescence, a striking decrease that correlated with the loss of pVCR94^GreenSp^. Conversely, low red fluorescence signal increased, consistent with chromosomally integrated SGI1^Red^. In contrast, when SGI1^Red^ Δ*sgaDC* was used, less than 4% of cells exhibited a high red fluorescence signal at T0, showing reduced SGI1 replication (Figure 7G). In addition, green and low red fluorescence remained steady throughout the experiment confirming that SGI1^Red^ remained chromosomally integrated and that pVCR94^GreenSp^ persisted in the cells. Remarkably, the presence of *acaDC* on the helper plasmid failed to trigger SGI1 replication in this context, confirming the importance of *sgaDC* in SGI1’s lifecycle. Complementation of Δ*sgaDC* using *pBAD-sgaDC* restored SGI1^Red^ replication and a strong incompatibility phenotype (Figure 6H). Remarkably, IncC plasmid loss correlated with decreased SGI1 replication and increased SGI1 integration, confirming that plasmid instability is caused by the rescue of *sgaDC* expression (Figure 6H). Finally, complementation using pBAD-*acaDC* triggered SGI1 excision but failed to restore SGI1 replication, resulting in progressive loss of both SGI1^Red^ and pVCR94^GreenSp^ (Figure 6I). Altogether, these data indicate that *sgaDC*, not *acaDC*, controls SGI1 excision and replication, and the subsequent destabilisation of the helper IncC plasmid.

### A single nucleotide polymorphism (SNP) abolishes SgaCD activity of SGI1-C

Contrasting with our observations with SGI1, deletion of *sgaDC* was shown to have no impact on the mobilisation of SGI1-C, whereas deletion of *acaDC* completely abolished both plasmid self-transfer and SGI1-C mobilisation (27, 28). Hence, *sgaDC* of SGI1-C is unable to complement an *acaDC* null mutant of its helper plasmid, unlike *sgaDC* of SGI1. To identify the cause of this discrepancy, we aligned *S008-sgaDC* genes and upstream sequences of SGI1 and SGI1-C and found only two SNPs in SGI1-C located at positions 6,600 (C to A) and 6,655 bp (G to A) relative to SGI1. While the second SNP is a silent mutation in *sgaC*, the first SNP changes the CTG codon to an ATG codon, resulting in a L139M substitution in SgaC C-terminal moiety immediately upstream of the predicted Zinc finger (Supplementary Figure S5F). A Blastp search of SgaC homologues revealed that either L or M amino acid residues are found at position 139. However, the M residue at this position is extremely rare in SGI1 variants (3 out the first 100 hits), with the notable exception of SGI2, formerly known as SGI1-J, of *S. enterica* serovar Emek from the UK (1999) (59). Furthermore, distantly related C subunits of other conjugative elements also have an L residue at the corresponding position (Supplementary Figure S5F), suggesting that the L139M substitution could be detrimental to their activity. FlhC, the most distant homologue, has a V residue at the corresponding position.

## DISCUSSION

SGI1 encodes the transcriptional activator complex SgaCD, whose biological relevance remained unclear until now (28). The similarity of SgaCD to other activator complexes, such as AcaCD encoded by IncC conjugative plasmids and SetCD encoded by SXT/R391 ICEs, suggested an important, yet perhaps redundant, role in SGI1 mobilisation (27, 34). Here we characterised the SgaCD regulon using ChIP-exo experiments, transcriptomic analyses, and β-galactosidase reporter assays. ChIP-exo experiments showed that SgaCD recognises and binds to the same sites as AcaCD both on the IncC plasmid and on SGI1, leading to activation of the transfer genes and operons (Figures 3 and 5). We also show that despite the evolutionary distance, both complexes bind the same DNA motifs and that *sgaDC* and *acaDC* are not exchangeable in their natural context, as a null mutant of either gene set exhibits drastically different phenotypes. While *acaDC* is essential to activate IncC plasmid transfer in the absence of SGI1, it becomes dispensable when SGI1 is present in the cell. In contrast, suppression of *sgaDC* of SGI1 in the presence of the IncC plasmid abolishes SGI1 replication and allows peaceful coexistence of both elements despite the presence of a fully functional copy of *acaDC*. Consistent with our results, SGI1-K that lacks *sgaDC* due to a 2,779-bp deletion extending from *S008* to the 5’ half of *traN_S_ (S005*) is compatible with IncC plasmids (33, 35). Furthermore, SGI1-C was reported to fail to complement an *acaDC* null mutant of its helper plasmid and the *sgaDC* deletion to have no impact on SGI1-C mobilisation (27, 28). We showed here that a single rare mutation in *sgaC* of SGI1-C, also found in SGI2, could be responsible for this phenotype, likely rendering SgaCD unable to act as a transcriptional activator at its physiological expression level.

The ability of SGI1 to complement an *acaDC* null IncC plasmid could have important epidemiological consequences. Occurrence of naturally *acaDC*-defective IncC plasmids has previously been reported (20). Although probably unable to activate self-transfer, entry of SGI1 in the host could resuscitate such “zombie” plasmids that would transiently regain their capacity to transfer, and mediate mobilisation of SGI1 (Supplementary Figure S6). This process would be facilitated by the ability of SGI1 to escape entry exclusion exerted by IncA and IncC plasmids that normally prevents or strongly reduces redundant transfer between cells that contain plasmids of the same entry exclusion group (31, 38).

The pathway allowing SGI1 to complement an *acaDC* null IncC plasmid is unclear, and could involve previously reported low-level, constitutive expression of *sgaDC* (27, 60). The absence of spontaneous excision of SGI1 suggests that the promoter of *sgaDC* is either mostly repressed under normal conditions or drastically activated by an IncA or IncC plasmid (61). Low SgaCD level produced by integrated SGI1 could be unable to switch on *xis* expression, preventing SGI1 excision and replication in the absence of an IncC plasmid. Likewise, low SgaCD level is probably insufficient to trigger expression of the entire IncC plasmid *tra* gene set and initiate transfer. However, since the rates of mobilisation of SGI1 and self-transfer of the Δ*acaDC* helper plasmid are comparable to wild-type (Figure 1A), we propose that an IncC plasmid-encoded factor, triggers a positive feed-back loop in response to low levels of SgaCD likely *via* activation of *sgaDC* expression. Alternatively, plasmid entry itself could eventually act as a trigger for activation of *sgaDC* expression (Figure 8), perhaps *via* activation of the SOS response by the invading single DNA strand during conjugation (62). Recent discovery of AcaB added an important piece to the regulatory switch that controls the “On/Off state” of conjugative transfer of IncA and IncC plasmids (25). Mutual activation of *acaB* and *acaDC* has been proposed to be the trigger or amplifier of the conjugative transfer “On state”. We showed here that, like AcaCD, SgaCD activates expression of AcaB (Figures 2 and 5), hence low level of SgaCD could initiate derepression of *P_acr1_ via* AcaB activation. This initial gentle push would then be amplified by AcaCD, promoting excision and replication of SGI1, which in return would raise SgaCD levels. However, although AcaB activates *acaDC* expression, it probably does not activate *sgaDC* expression, since no AcaCD- or AcaB-binding site was found upstream of *sgaDC* (20, 25). Yet another IncC-encoded factor is likely at play. Consistent with this hypothesis, SGI1’s *rep* gene was shown to be slightly expressed in the presence of pVCR94^Sp^ Δ*acaDC*, but not in its absence (30). Since we showed here that deletion of *sgaDC* inhibits the excision of SGI1, abolishes its replication, reduces considerably its rate of mobilisation and alleviates incompatibility with IncC plasmids (Figure 1A), this observation supports the existence of a plasmid-encoded activator of *sgaDC* expression. Hence, the promoter of *sgaDC* instead of *P_xis_* as previously suggested (27) would act as the sensor for IncC plasmid entry. Identification of the putative factor promoting such a feedback loop is on-going.

**Figure 8.**
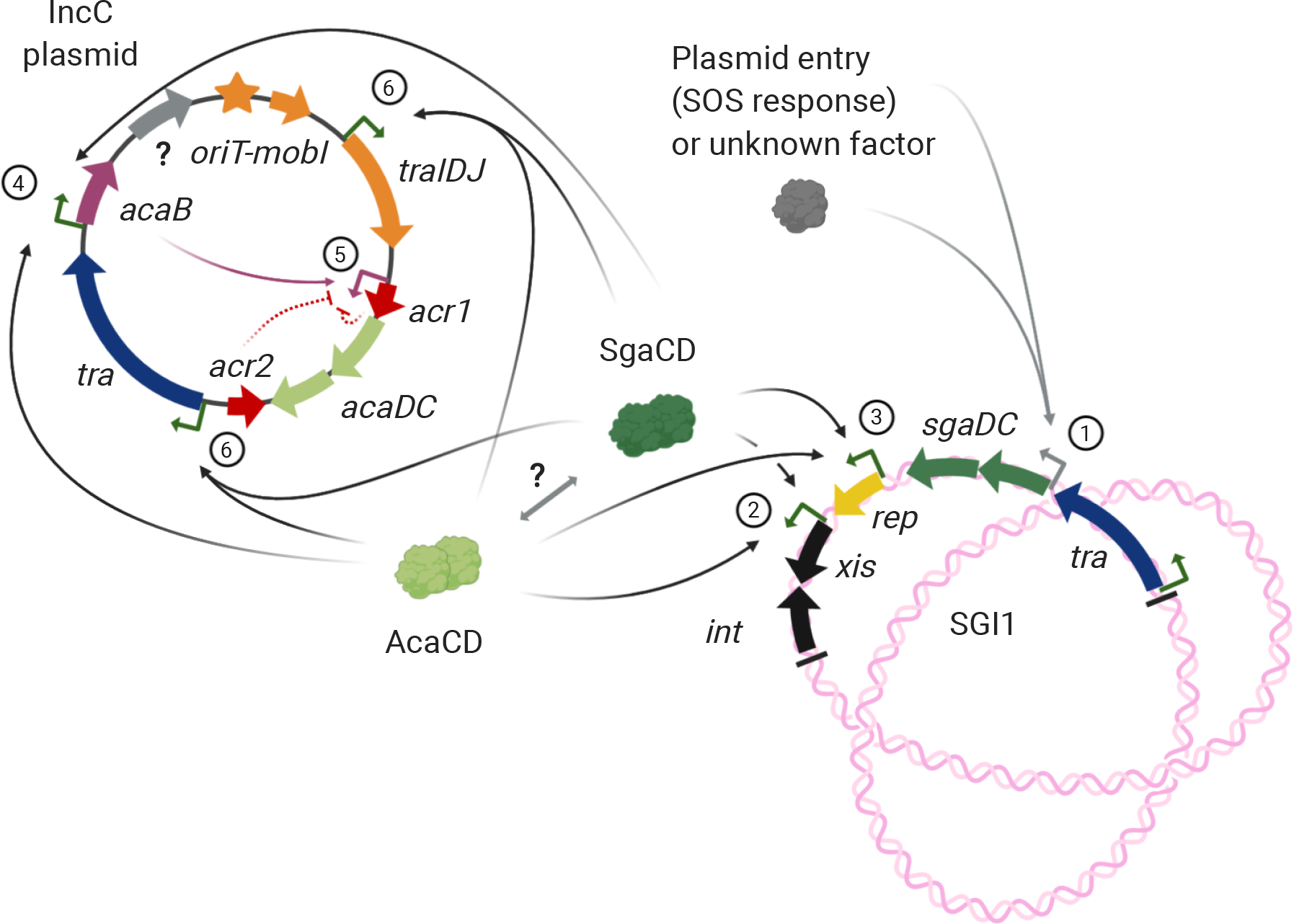
Model of regulation of IncC plasmids and SGI1 gene expression. Major genes and operons involved in conjugative transfer and regulation are depicted as color-coded arrowed boxes (or a star for IncC origin of transfer *oriT*) based on the function: green, purple or grey, transcriptional activation; red, transcriptional repression; blue, type IV secretion system; orange, relaxosome; black, site-specific recombination; yellow, replication. Promoters are depicted by angled arrows and color-coded based on the corresponding activator: grey, unknown; green, AcaCD/SgaCD-activatable; purple, AcaB-activatable. Activation is represented by faded arrows. Repression is represented by red blocked dashed arrows. A bidirectional grey arrow indicates potential interactions. The chromosome is depicted as a large knotted DNA structure. SGI1 is represented in its initial integrated form, with its attachment sites depicted as two black lines. We propose a model in 6 main steps: 1) activation of *sgaDC* expression, 2) activation of SGI1 excision, 3) activation of SGI1 replication, 4) activation of *acaB*, 5) activation of *acaDC* and 6) activation of all the AcaCD/SgaCD-dependent promoters. For clarity, the activation of only two IncC transfer operons is depicted. Timing of the alleviation of repression by Acr1 and Acr2 is unknown. Created with BioRender.com.

By analogy with the flagellar activator FlhCD to which they are distantly related, the C subunit of AcaCD and SgaCD likely binds to DNA immediately upstream of the −35 element, whereas the D subunit would stabilise the transcriptional complex (63). In support of this hypothesis, the primary sequence of D subunits is more divergent than the one of C subunits (27). Furthermore, AcaC and SgaC also share predicted tertiary structures that are remarkably similar to that of FlhC, including the Zinc-finger domain and its four conserved cysteine residues (Supplementary Figure S5). The similarity of SgaC and AcaC likely accounts for the cross recognition of the same set of binding sites. Considering that both complexes derived from a common ancestor, weaker promoter activation by SgaCD, together with switchable *sgaDC* expression, could have evolved during its domestication by SGI1-like elements to reduce the risk of futile excision in the absence of a helper plasmid, enhancing SGI1 stability and is compensated by its ability to replicate. The IncC plasmid maintains as a single-copy replicon per cell, whereas the replicative cycle of SGI1 in IncC^+^ cells generates over 7 copies per cell, likely enhancing *sgaDC* expression (Figure 6B) (30). Several properties of the expression of the transcriptional activator genes remain to be characterised to fully understand the crosstalk between SGI1 and its helper plasmid, including the conditions of activation and strength of *sgaDC* and *acaDC* promoters, the halflife of the corresponding mRNA transcripts, their translation rates and the relative stability of each activator complex. Assuming that SgaCD is produced in a much larger quantity than AcaCD due to SGI1’s high copy number, with comparable turnover rates of both complexes, AcaCD’s role could become negligible and even dispensable. This hypothesis is supported not only by the full complementation of an *acaDC* null mutant by SGI1, but also by the significant increase of plasmid transfer upon *sgaDC* expression (Figure 1). We cannot rule out that chimerical activator complexes SgaC-AcaD or AcaC-SgaD could form and play an important role in the regulation of gene activation on the helper plasmid and on SGI1. If such chimera exist, and if SgaCD and AcaCD tend to form heterohexamers like *E. coli* FlhD_4_C_2_ (63), then a heterogeneous bestiary of activator complexes regulates gene expression when SGI1 and its helper plasmid occupy the same cell.

Incompatibility between SGI1 and IncC plasmids could have emerged as a defence mechanism deployed by SGI1 to prevent excessive activation of excision and replication, which ultimately results in SGI1 loss in the absence of selective pressure (Figure 7). Destabilisation of the plasmid would allow the island to revert to its quiescent state after transferring to a new host, ensuring its stability by residing integrated in the host’s chromosome. Although the exact mechanism lying underneath the destabilisation is still unknown, we recently suggested titration of endogenous replication proteins (30). Alternatively, SGI1 could be interfering with the partitioning of the IncC plasmid in daughter cells. Being single-copy large plasmids (Figure 6C), IncC replicons are likely extremely susceptible to perturbation of their partition process. In fact, IncC plasmids encode two partitioning systems, a type I *parABS* that has been shown to be essential for plasmid maintenance and a putative type II *parMRC-like* partitioning system distantly related to *srpRMC* of SXT/R391 ICEs and encoded by *vcrx151-152* (64, 65). We show here that expression of *vcrx151-152* is activated by AcaCD and SgaCD. Remarkably, forced excision and replication of SGI1 *via* overexpression of *acaDC* or *sgaDC* in the absence of an IncC plasmid resulted in rapid elimination of SGI1 from the cell population (Figure 7D-E). In stark contrast, when the plasmid was present, SGI1 persisted and reintegrated in the chromosome while promoting IncC plasmid loss (Figure 7H). This observation hints at SGI1 taking control of the IncC partitioning systems perhaps to enhance its equal segregation into daughter cells during cell division. These lines of inquiry will need to be examined in detail to better understand the processes at stake.

Despite the divergence of the primary sequences of AcaC and SgaC, we showed that both transcriptional activator complexes bind to the same DNA motif. Other conjugative plasmids that confer multidrug resistance to marine-dwelling bacteria and have not yet been ascribed to an incompatibility group also encode AcaCD orthologues. The C-terminus of the C subunits encoded by these plasmids (e.g. AqaC or AsaC) diverge significantly from AcaC and SgaC. Nevertheless, their Zinc-finger domain is relatively well conserved (Supplementary Figure 5F-G). Unsurprisingly, these plasmids contain AcaCD-like binding sites upstream genes involved in conjugative transfer (Supplementary Table S4). Based on these observations, the breadth of IncC/SGI1-like interactions is likely broader than anticipated, with SGI1 and its variants being possibly activated and mobilized by such plasmids that do not belong to the IncA nor the IncC group (16). Furthermore, since AcaCD-responsive promoters have recently been found in MGIs integrated at *trmE, yicC* and *dusA* in the chromosome of several species of Gammaproteobacteria (9), our observations support a complex network of mobilisation events involving diverse families of MGIs and conjugative plasmids of multiple incompatibility groups (IncA, IncC and untyped) encoding AcaCD-like transcriptional activators with identical an DNA-binding motif.

Interactions between members of the SGI1 family and their helper plasmids are undoubtedly complex. The use of naturally occurring variants such as SGI1-C, SGI1-F, SGI1-I, SGI1-K or SGI2, with diverse naturally occurring IncC and even IncA plasmids, while providing interesting hints and clues, also renders comparisons between studies challenging and could lead to erroneous conclusions. These shortcomings result from SNPs that may affect the expression of key effectors that are important for the regulation, replication, stability, and conjugative transfer of the two interacting partners. Considering the discrepancies between others’ results and ours, we urge the establishment of a reliable, robust system based on a selected subset of model SGI1 variants and helper plasmids to properly decipher the complex biology and interactions between SGI1 and IncC plasmids.

## AVAILABILITY

Complete data from aligned reads for ChIP-Exo, Cappable-Seq and RNA-Seq experiments can also be visualized using the UCSC genome browser at http://bioinfo.ccs.usherbrooke.ca/sgaCD.html

## ACCESSION NUMBERS

Raw sequencing data were submitted to Genbank under Bioproject accession number PRJNA648047 with the following Biosample accession numbers: for ChIP-exo assays, from SAMN15617565 to SAMN15617572, respectively; for Cappable-Seq assays, from SAMN15617573 to SAMN15617580; for RNA-Seq assays, from SAMN15617581 to SAMN15617604.

## SUPPLEMENTARY DATA

Supplementary Data are available at NAR online.

## ACKNOWLEDGEMENT

We thank Calcul Quebec (www.calculquebec.ca) and Compute Canada (www.computecanada.ca) for access to bioinformatics resources and support, as well as Jean-François Lucier for his help with bioinformatic analyses. We also thank the Centre de calcul scientifique de l’Université de Sherbrooke for technical assistance and the Université de Sherbrooke RNomics Plateform for assistance with Illumina sequencing. We are grateful to Alain Lavigueur for his insightful comments on the manuscript and to Dominick Matteau for thoughtful discussions and advice.

## FUNDING

This work was supported by a Discovery Grant [2016-04365] from the Natural Sciences and Engineering Research Council of Canada (NSERC) and a Project Grant [PJT-153071] from the Canadian Institutes of Health Research (CIHR) to V.B. R.D. is the recipient of a Fonds de recherche du Québec-Nature et Technologies (FRQNT) doctoral fellowship K.T.H. was supported by a postdoctoral fellowship [SPE20170336797] from the Fondation de la Recherche Médicale (FRM, France). N.R. is the recipient of a Fonds de recherche du Québec-Santé (FRQS) MSc scholarship and an Alexander Graham Bell Canada Graduate Scholarship from the NSERC. Funding for open access charge: NSERC [2016-04365].

## CONFLICT OF INTEREST

The authors declare no competing interests.

## Notes

http://bioinfo.ccs.usherbrooke.ca/sgaCD.html

### Competing Interest Statement

The authors have declared no competing interest.

## REFERENCES

1. CDC (2019) Antibiotic Resistance Threats in the United States, 2019. Atlanta, GA: U.S. Department of Health and Human Services, CDC.

2. MacLean, R.C. and Millan, A.S. (2019) The evolution of antibiotic resistance. Science, 365, 1082–1083.

3. Guédon, G., Libante, V., Coluzzi, C., Payot, S. and Leblond-Bourget, N. (2017) The Obscure World of Integrative and Mobilizable Elements, Highly Widespread Elements that Pirate Bacterial Conjugative Systems. Genes, 8, 337.

4. Bellanger, X., Payot, S., Leblond-Bourget, N. and Guédon, G. (2014) Conjugative and mobilizable genomic islands in bacteria: evolution and diversity. FEMS Microbiology Reviews, 38, 720–760.

5. Carraro, N., Rivard, N., Burrus, V. and Ceccarelli, D. (2017) Mobilizable genomic islands, different strategies for the dissemination of multidrug resistance and other adaptive traits. Mob Genet Elements, 7, 1–6.

6. Piña-Iturbe, A., Ulloa-Allendes, D., Pardo-Roa, C., Coronado-Arrázola, I., Salazar-Echegarai, F.J., Sclavi, B., González, P.A. and Bueno, S.M. (2018) Comparative and phylogenetic analysis of a novel family of Enterobacteriaceae-associated genomic islands that share a conserved excision/integration module. Sci Rep, 8, 10292.

7. Carraro, N., Rivard, N., Ceccarelli, D., Colwell, R.R. and Burrus, V. (2016) IncA/C conjugative plasmids mobilize a new family of multidrug resistance islands in clinical *Vibrio cholerae* non-O1/non-O139 isolates from Haiti. mBio, 7, pii: e00509–16.

8. Brouwer, M.S.M., Warburton, P.J., Roberts, A.P., Mullany, P. and Allan, E. (2011) Genetic organisation, mobility and predicted functions of genes on integrated, mobile genetic elements in sequenced strains of *Clostridium difficile*. PLoS ONE, 6, e23014.

9. Rivard, N., Colwell, R.R. and Burrus, V. (2020) Antibiotic Resistance in *Vibrio cholerae:* Mechanistic Insights from IncC Plasmid-Mediated Dissemination of a Novel Family of Genomic Islands Inserted at *trmE*. mSphere, 5, e00748–20.

10. Boyd, D., Peters, G.A., Cloeckaert, A., Boumedine, K.S., Chaslus-Dancla, E., Imberechts, H. and Mulvey, M.R. (2001) Complete nucleotide sequence of a 43-kilobase genomic island associated with the multidrug resistance region of *Salmonella enterica* serovar Typhimurium DT104 and its identification in phage type DT120 and serovar Agona. J. Bacteriol., 183, 5725–5732.

11. Cummins, M.L., Hamidian, M. and Djordjevic, S.P. (2020) *Salmonella* Genomic Island 1 is Broadly Disseminated within Gammaproteobacteriaceae. Microorganisms, 8, 161.

12. Schultz, E., Barraud, O., Madec, J.-Y., Haenni, M., Cloeckaert, A., Ploy, M.-C. and Doublet, B. (2017) Multidrug Resistance *Salmonella* Genomic Island 1 in a *Morganella morganii* subsp. *morganii* Human Clinical Isolate from France. mSphere, 2, e00118–17.

13. Soliman, A.M., Shimamoto, T., Nariya, H. and Shimamoto, T. (2019) Emergence of *Salmonella* Genomic Island 1 Variant SGI1-W in a Clinical Isolate of *Providencia stuartii* from Egypt. Antimicrob. Agents Chemother., 63, e01793–18.

14. Siebor, E. and Neuwirth, C. (2013) Emergence of *Salmonella* genomic island 1 (SGI1) among *Proteus mirabilis* clinical isolates in Dijon, France. J. Antimicrob. Chemother., 68,1750–1756.

15. Soliman, A.M., Ramadan, H., Ghazy, E., Yu, L., Hisatsune, J., Kayama, S., Sugai, M., Nariya, H., Shimamoto, T., Jackson, C.R., et al. (2020) Emergence of Salmonella genomic island 1 variant SGI1-C in a multidrug-resistant clinical isolate of *Klebsiella pneumoniae* ST485 from Egypt. Antimicrob. Agents Chemother., 64, e01055–20.

16. Douard, G., Praud, K., Cloeckaert, A. and Doublet, B. (2010) The *Salmonella* genomic island 1 is specifically mobilized in trans by the IncA/C multidrug resistance plasmid family. PLoS ONE, 5, e15302.

17. Wu, W., Feng, Y., Tang, G., Qiao, F., McNally, A. and Zong, Z. (2019) NDM Metallo-β-Lactamases and Their Bacterial Producers in Health Care Settings. Clin. Microbiol. Rev., 32, e00115–18.

18. Weill, F.-X., Domman, D., Njamkepo, E., Tarr, C., Rauzier, J., Fawal, N., Keddy, K.H., Salje, H., Moore, S., Mukhopadhyay, A.K., et al. (2017) Genomic history of the seventh pandemic of cholera in Africa. Science, 358, 785–789.

19. Arcari, G., Di Lella, F.M., Bibbolino, G., Mengoni, F., Beccaccioli, M., Antonelli, G., Faino, L. and Carattoli, A. (2020) A Multispecies Cluster of VIM-1 Carbapenemase-Producing *Enterobacterales* Linked by a Novel, Highly Conjugative, and Broad-Host-Range IncA Plasmid Forebodes the Reemergence of VIM-1. Antimicrob Agents Chemother, 64, e02435–19, /aac/64/4/AAC.02435-19.atom.

20. Carraro, N., Matteau, D., Luo, P., Rodrigue, S. and Burrus, V. (2014) The master activator of IncA/C conjugative plasmids stimulates genomic islands and multidrug resistance dissemination. PLoS Genet, 10, e1004714.

21. Poulin-Laprade, D., Matteau, D., Jacques, P.-É., Rodrigue, S. and Burrus, V. (2015) Transfer activation of SXT/R391 integrative and conjugative elements: unraveling the SetCD regulon. Nucleic Acids Res., 43, 2045–2056.

22. Liu, X. and Matsumura, P. (1994) The FlhD/FlhC complex, a transcriptional activator of the *Escherichia coli* flagellar class II operons. J. Bacteriol., 176, 7345–7351.

23. Beaber, J.W., Hochhut, B. and Waldor, M.K. (2002) Genomic and functional analyses of SXT, an integrating antibiotic resistance gene transfer element derived from *Vibrio cholerae*. J. Bacteriol., 184, 4259–4269.

24. Lang, K.S. and Johnson, T.J. (2016) Characterization of Acr2, an H-NS-like protein encoded on A/C_2_-type plasmids. Plasmid, 87–88, 17–27.

25. Hancock, S.J., Phan, M.-D., Luo, Z., Lo, A.W., Peters, K.M., Nhu, N.T.K., Forde, B.M., Whitfield, J., Yang, J., Strugnell, R.A., et al. (2020) Comprehensive analysis of IncC plasmid conjugation identifies a crucial role for the transcriptional regulator AcaB. Nat Microbiol, 10.1038/s41564-020-0775-0.

26. Carraro, N., Matteau, D., Burrus, V. and Rodrigue, S. (2015) Unraveling the regulatory network of IncA/C plasmid mobilization: When genomic islands hijack conjugative elements. Mob Genet Elements, 5, 1–5.

27. Kiss, J., Papp, P.P., Szabó, M., Farkas, T., Murányi, G., Szakállas, E. and Olasz, F. (2015) The master regulator of IncA/C plasmids is recognized by the *Salmonella* Genomic island SGI1 as a signal for excision and conjugal transfer. Nucleic Acids Res., 43, 8735–8745.

28. Murányi, G., Szabó, M., Olasz, F. and Kiss, J. (2016) Determination and Analysis of the Putative AcaCD-Responsive Promoters of *Salmonella* Genomic Island 1. PLoS ONE, 11, e0164561.

29. Doublet, B., Boyd, D., Mulvey, M.R. and Cloeckaert, A. (2005) The *Salmonella* genomic island 1 is an integrative mobilizable element. Mol. Microbiol., 55, 1911–1924.

30. Huguet, K.T., Rivard, N., Garneau, D., Palanee, J. and Burrus, V. (2020) Replication of the *Salmonella* Genomic Island 1 (SGI1) triggered by helper IncC conjugative plasmids promotes incompatibility and plasmid loss. PLoS Genet., 16, e1008965.

31. Carraro, N., Durand, R., Rivard, N., Anquetil, C., Barrette, C., Humbert, M. and Burrus, V. (2017) *Salmonella* genomic island 1 (SGI1) reshapes the mating apparatus of IncC conjugative plasmids to promote self-propagation. PLOS Genetics, 13, e1006705.

32. Huguet, K.T., Gonnet, M., Doublet, B. and Cloeckaert, A. (2016) A toxin antitoxin system promotes the maintenance of the IncA/C-mobilizable *Salmonella* Genomic Island 1. Sci Rep, 6, 32285.

33. Harmer, C.J., Hamidian, M., Ambrose, S.J. and Hall, R.M. (2016) Destabilization of IncA and IncC plasmids by SGI1 and SGI2 type *Salmonella* genomic islands. Plasmid, 87–88, 51–57.

34. Poulin-Laprade, D., Carraro, N. and Burrus, V. (2015) The extended regulatory networks of SXT/R391 integrative and conjugative elements and IncA/C conjugative plasmids. Front Microbiol, 6, 837.

35. Hamidian, M., Holt, K.E. and Hall, R.M. (2015) The complete sequence of *Salmonella* genomic island SGI1-K. J. Antimicrob. Chemother., 70, 305–306.

36. Boyd, D., Cloeckaert, A., Chaslus-Dancla, E. and Mulvey, M.R. (2002) Characterization of variant *Salmonella* genomic island 1 multidrug resistance regions from serovars Typhimurium DT104 and Agona. Antimicrob. Agents Chemother., 46, 1714–1722.

37. Dower, W.J., Miller, J.F. and Ragsdale, C.W. (1988) High efficiency transformation of *E. coli* by high voltage electroporation. Nucleic Acids Res., 16, 6127–6145.

38. Humbert, M., Huguet, K.T., Coulombe, F. and Burrus, V. (2019) Entry Exclusion of Conjugative Plasmids of the IncA, IncC, and Related Untyped Incompatibility Groups. J. Bacteriol., 201, e00731–18.

39. Datsenko, K.A. and Wanner, B.L. (2000) One-step inactivation of chromosomal genes in *Escherichia coli* K-12 using PCR products. Proc Natl Acad Sci U S A, 97, 6640–6645.

40. Haldimann, A. and Wanner, B.L. (2001) Conditional-replication, integration, excision, and retrieval plasmid-host systems for gene structure-function studies of bacteria. J. Bacteriol., 183, 6384–6393.

41. Ettwiller, L., Buswell, J., Yigit, E. and Schildkraut, I. (2016) A novel enrichment strategy reveals unprecedented number of novel transcription start sites at single base resolution in a model prokaryote and the gut microbiome. BMC Genomics, 17, 199.

42. Christodoulou, D.C., Gorham, J.M., Herman, D.S. and Seidman, J.G. (2011) Construction of normalized RNA-seq libraries for next-generation sequencing using the crab duplex-specific nuclease. Curr Protoc Mol Biol, Chapter 4, Unit 4.12.

43. Bolger, A.M., Lohse, M. and Usadel, B. (2014) Trimmomatic: a flexible trimmer for Illumina sequence data. Bioinformatics, 30, 2114–2120.

44. Andrews, S. (2010) FastQC: A Quality Control tool for High Throughput Sequence Data.

45. Langmead, B. and Salzberg, S.L. (2012) Fast gapped-read alignment with Bowtie 2. Nat Methods, 9, 357–359.

46. Lassmann, T., Hayashizaki, Y. and Daub, C.O. (2011) SAMStat: monitoring biases in next generation sequencing data. Bioinformatics, 27, 130–131.

47. Li, H., Handsaker, B., Wysoker, A., Fennell, T., Ruan, J., Homer, N., Marth, G., Abecasis, G. and Durbin, R. (2009) The Sequence Alignment/Map format and SAMtools. Bioinformatics, 25, 2078–2079.

48. Quinlan, A.R. and Hall, I.M. (2010) BEDTools: a flexible suite of utilities for comparing genomic features. Bioinformatics, 26, 841–842.

49. Coulombe, C., Poitras, C., Nordell-Markovits, A., Brunelle, M., Lavoie, M.-A., Robert, F. and Jacques, P.-É. (2014) VAP: a versatile aggregate profiler for efficient genome-wide data representation and discovery. Nucleic Acids Res., 42, W485–493.

50. Magoc, T., Wood, D. and Salzberg, S.L. (2013) EDGE-pro: Estimated Degree of Gene Expression in Prokaryotic Genomes. Evol Bioinform Online, 9, 127–136.

51. Love, M.I., Huber, W. and Anders, S. (2014) Moderated estimation of fold change and dispersion for RNA-seq data with DESeq2. Genome Biology, 15, 550.

52. Altschul, S.F., Gish, W., Miller, W., Myers, E.W. and Lipman, D.J. (1990) Basic local alignment search tool. J. Mol. Biol., 215, 403–410.

53. Miller, J.H. (1992) A Short Course in Bacterial Genetics: A Laboratory Manual and Handbook for *Escherichia coli* and Related Bacteria Cold Spring Harbor Laboratory Press, Cold Spring Harbor.

54. Hellemans, J., Mortier, G., De Paepe, A., Speleman, F. and Vandesompele, J. (2007) qBase relative quantification framework and software for management and automated analysis of real-time quantitative PCR data. Genome Biology, 8, R19.

55. Browning, D.F. and Busby, S.J. (2004) The regulation of bacterial transcription initiation. Nat. Rev. Microbiol., 2, 57–65.

56. Latif, H., Federowicz, S., Ebrahim, A., Tarasova, J., Szubin, R., Utrilla, J., Zengler, K. and Palsson, B.O. (2018) ChIP-exo interrogation of Crp, DNA, and RNAP holoenzyme interactions. PLoS ONE, 13, e0197272.

57. Hegyi, A., Szabó, M., Olasz, F. and Kiss, J. (2017) Identification of *oriT* and a recombination hot spot in the IncA/C plasmid backbone. Sci Rep, 7, 10595.

58. Carraro, N., Sauvé, M., Matteau, D., Lauzon, G., Rodrigue, S. and Burrus, V. (2014) Development of pVCR94ΔX from *Vibrio cholerae*, a prototype for studying multidrug resistant IncA/C conjugative plasmids. Front Microbiol, 5, 44.

59. Hamidian, M., Holt, K.E. and Hall, R.M. (2015) The complete sequence of *Salmonella* genomic island SGI2. Journal of Antimicrobial Chemotherapy, 70, 617–619.

60. Golding, G.R., Olson, A.B., Doublet, B., Cloeckaert, A., Christianson, S., Graham, M.R. and Mulvey, M.R. (2007) The effect of the *Salmonella* genomic island 1 on in vitro global gene expression in *Salmonella enterica* serovar Typhimurium LT2. Microbes Infect., 9, 21–27.

61. Kiss, J., Nagy, B. and Olasz, F. (2012) Stability, entrapment and variant formation of *Salmonella* genomic island 1. PLoS ONE, 7, e32497.

62. Baharoglu, Z., Bikard, D. and Mazel, D. (2010) Conjugative DNA transfer induces the bacterial SOS response and promotes antibiotic resistance development through integron activation. PLoS Genet., 6, e1001165.

63. Wang, S., Fleming, R.T., Westbrook, E.M., Matsumura, P. and McKay, D.B. (2006) Structure of the *Escherichia coli* FlhDC complex, a prokaryotic heteromeric regulator of transcription. J. Mol. Biol., 355, 798–808.

64. Carraro, N., Poulin, D. and Burrus, V. (2015) Replication and Active Partition of Integrative and Conjugative Elements (ICEs) of the SXT/R391 Family: The Line between ICEs and Conjugative Plasmids Is Getting Thinner. PLoS Genet, 11, e1005298.

65. Hancock, S.J., Phan, M.-D., Peters, K.M., Forde, B.M., Chong, T.M., Yin, W.-F., Chan, K.-G., Paterson, D.L., Walsh, T.R., Beatson, S.A., et al. (2017) Identification of IncA/C Plasmid Replication and Maintenance Genes and Development of a Plasmid Multilocus Sequence Typing Scheme. Antimicrob. Agents Chemother., 61, e01740–16.

66. Grenier, F., Matteau, D., Baby, V. and Rodrigue, S. (2014) Complete genome sequence of *Escherichia coli* BW25113. Genome Announc, 2, pii: e01038–14.

67. Singer, M., Baker, T.A., Schnitzler, G., Deischel, S.M., Goel, M., Dove, W., Jaacks, K.J., Grossman, A.D., Erickson, J.W. and Gross, C.A. (1989) A collection of strains containing genetically linked alternating antibiotic resistance elements for genetic mapping of Escherichia coli. Microbiol Rev, 53, 1–24.

68. Ceccarelli, D., Daccord, A., René, M. and Burrus, V. (2008) Identification of the Origin of Transfer *(oriT*) and a New Gene Required for Mobilization of the SXT/R391 Family of Integrating Conjugative Elements. J. Bacteriol., 190, 5328–5338.

69. Datta, S., Costantino, N. and Court, D.L. (2006) A set of recombineering plasmids for gram-negative bacteria. Gene, 379, 109–115.

70. Cherepanov, P.P. and Wackernagel, W. (1995) Gene disruption in *Escherichia coli:* TcR and KmR cassettes with the option of Flp-catalyzed excision of the antibioticresistance determinant. Gene, 158, 9–14.

71. Guzman, L.M., Belin, D., Carson, M.J. and Beckwith, J. (1995) Tight regulation, modulation, and high-level expression by vectors containing the arabinose *P_BAD_* promoter. J. Bacteriol., 177, 4121–4130.

