## Supplementary data for "Crucial Role of *Salmonella* Genomic Island 1 Master Activator in Parasitism of IncC plasmids"

**Supplementary Table 1. Oligonucleotides used in this study.**

| Primer | Sequence | Reference |
|---|---|---|
| SGI1delxis.for | AACTAGGTCTATCAATAACTGGTGTGTAACGGGTAGGCTGgtgtaggctggagctgcttc | This study |
| SGI1delxis.rev | CTCACTGCGGAAAGTCCGTATTGAATAAAACACTTTAAGAttattaCATATGAATATCCTCCTTA | This study |
| SGI1delsgaC06.for | TATTAAAAAAGATTCAGCAATCTTTGGTGTGGAGATAATCgtgtaggctggagctgcttc | This study |
| SGI1delsgaC06.rev | GCTATTTGCCCTTTTGCGGCATACGCGGATGTATTCAttaCATATGAATATCCTCCTTA | This study |
| SGI1delsgaD07.for | CAATCTGGGCCGCAGAAAAAAAGGTAAGGAGGACTACTGAgtgtaggctggagctgcttc | This study |
| SGI1delsgaD07.rev | TTACCCAGCGAGATAACGAACCAGATGTAAAACATTCACTcatatgaatatcctcctta | This study |
| pVCR94delY2.f | TTTAATATTTACAGCATCACATAATGATACTATGATGTTGcggaataggaacttcaagaat | This study |
| 94DelXnoFRT.rev | TCGTTAACTGCACATTCGGGATATTTCTCTATATTCGCGGagagcgcttttgaagctca | (1) |
| 94DelacaD.for | GCTATCTATCGCAACCTTCGTGATTTGTGAGGGGGGCGGAgtgtaggctggagctgcttc | (2) |
| 94DelacaC.rev | GCGCTCATCTTCTGGTCCGAAATGTCATAGTCTACTCAttaCATATGAATATCCTCCTTA | (2) |
| SGI1sgaDEcoRI.for | NNNNNNGAATTCaaggaggaataataaATGTATGCCTTAGAGCCGTT | This study |
| SGI1sgaCEcoRI.rev | NNNNNNGAATTCTCAGGCAGCTTTTGAGA | This study |
| pAH56sgaCDvecF | aagctgccAAGCTTGACTACAAAGAC | This study |
| pAH56sgaCDvecR | taaggcataCATATGTATATCTCCTTCTTACAAG | This study |
| pAH56sgaCDinsF | tatacatatgTATGCCTTAGAGCCGTTAGAAC | This study |
| pAH56sgaCDinsR | gtcaagcttGGCAGCTTTTGAGACCCG | This study |
| promvcrx012PstIF | NNCTGCAGGTGGCGGAAAGTTTCACAT | This study |
| promvcrx012XhoIR | NNNNCTCGAGTATTCACCTCCTATACCAACCAG | This study |
| promvcrx36PstIF | NNCTGCAGAACAAAGCTCCTTAATTGCTTG | This study |
| promvcrx35PstIF | NNCTGCAGACATATCACCTTTACAGTATCATT | This study |
| promvcrx060traIpstI.for | NNNNNNCTGCAGCATCAAAAATTGTCGATGA | (2) |
| promvcrx060traIpstI.rev | NNNNNNCTGCAGCTATCGTATTTCTCGTCGCTA | (2) |
| promvcrx059XhoIR | NNNNCTCGAGCATCAAAAATTGTCGATGA | This study |
| promvcrx068PstIF | NNCTGCAGCCTCCATAAAGTGCCCAAAATG | This study |
| promvcrx068XhoIR | NNNNCTCGAGAGACCCTCCAAATAACTTGTCGT | This study |
| promtraLpstI.for | NNNNNNCTGCAGCCGATCCAGTAAACGCAA | (2) |
| promtraLpstI.rev | NNNNNNCTGCAGTTCTTTTCCTAGTGGCCTGTA | (2) |
| promtraVPstIF | NNCTGCAGTGTTCTCCGACCAGATGTTG | This study |
| promtraAXhoIR | NNNNCTCGAGCCGACAAAATAAGTAGCGCTG | This study |
| promvcrx076PstIF | NNCTGCAGAAGACACCTTGCCGTT | This study |
| promvcrx076XhoIR | NNNNCTCGAGCGCATTCTCCATTCGTTTTG | This study |
| promdsbCPstIF | NNCTGCAGGGTTCATTCCTCTCAAAGC | This study |

| | | |
|---|---|---|
| promdsbCXhoIR | NNNNCTCGAGGTCTATCCCCTAAGATTGGATTA | This study |
| promtraNcPstIF | NNCTGCAGGCAGACCCTAACGAATTTG | This study |
| promtraNcXhoIR | NNNNCTCGAGTGTGTTCTCCATTTCAACCTT | This study |
| promvcrx087PstIF | NNCTGCAGTAGAGATACTCCTAAGTTGAGGTA | This study |
| promvcrx086PstIF | NNCTGCAGGGTGTTGCTCCTATAAGAA | This study |
| promvcrx098PstIF | NNCTGCAGTGCGTCTTTGAGCTTTG | This study |
| promvcrx098XhoIR | NNNNCTCGAGGTGTGCTCCTTAAAGTAACTTG | This study |
| promvcrx114PstIF | NNCTGCAGAAGGCCCTCCTTCG | This study |
| promvcrx114XhoIR | NNNNCTCGAGGCTTCTCTCCTTTGTTTGTG | This study |
| promvcrx128PstIFv2 | NNCTGCAGGTAACCTGCTTCTCATTTGTTCTG | This study |
| promvcrx128XhoIR | NNNNCTCGAGGAAGGAATGCGCGTCTATG | This study |
| promvcrx140PstIF | NNCTGCAGCAGGATCACCTGGAGCAAG | This study |
| promvcrx140XhoIR | NNNNCTCGAGGAGCTGCTCTTTCAACGATG | This study |
| promtraFpstI.for | NNNNNNCTGCAGGTTTGGTCAATTCTTTCACAT | (2) |
| promtraFpstI.rev | NNNNNNCTGCAGCCATGCTCGATATGGTAGA | (2) |
| promacaQpstI.for | NNNNNNCTGCAGTACTCTTTACCTCCAGTTTACCA | (2) |
| promacaQpstI.rev | NNNNNNCTGCAGTTAAACTGCGTTGTTAGCCAT | (2) |
| promvcrx152/001pstI.for | NNNNNNCTGCAGCAAAGATTGCTTTTAGATTGCTT | (2) |
| promvcrx152/001pstI.rev | NNNNNNCTGCAGCAATTCCTATCCAATTCCTA | (2) |
| SGI1promxisPstI.for | NNNNNNCTGCAGGTTATTGATAGACCTAGTTTAT | (3) |
| SGI1promxisPstI.rev | NNNNNNCTGCAGGGCCAATGTGCCGGTTT | (3) |
| SGI1promS004PstI.for | NNNNNNCTGCAGGGCCTCTGGAAGTGCCCTAT | This study |
| proms004XhoIR | NNNNCTCGAGTAAATTTCTCCAGCATCATCATTGATTTACTG | This study |
| SGI1promtraNPstI.for | NNNNNNCTGCAGCCAGCTTTTTAGTTTGGATA | (3) |
| SGI1promtraNPstI.rev | NNNNNNCTGCAGGATAGAGCATTGCGAGCAAT | (3) |
| SGI1promtraHPstI.for | NNNNNNCTGCAGTAGGTTTCGTGTCACCCGAA | (3) |
| SGI1promtraHPstI.rev | NNNNNNCTGCAGTTAAAGCTCCTCTTTTAGAA | (3) |
| SGI1proms018PstI.for | NNNNNNCTGCAGAATTAGTTGGAATTGAAGGGGTT | This study |
| SGI1proms018PstI.rev | NNNNNNCTGCAGGGCTCTGCTGATTAAAAGCCAT | This study |
| qAttBFw | AACATCTACAACAGGGCAAAG | (4) |
| qAttBRv | GAGGAATAACAGGAGTGGTAAC | (4) |
| qS026Fw | TGTCATCAGAAAGAACAAGCTC | (4) |
| qS026Rv | GCGTTTTTATTCTGTTGCCC | (4) |
| qFwpVCR | AAGAGAACCAAAGACAAAGACC | (4) |
| qRvpVCR | CACCTTCACCGTGAAATGC | (4) |
| qdnaBFw | ACGATTTTTACACCCGCCCAC | (4) |
| qdnaBRv | ATCATCTCACGGACAACGGCAC | (4) |
| qhicBFw | GCTTATCCCTTTACCTTCGCC | (4) |
| qhicBRv | TAACTCTTTGCCAAGCGCC | (4) |

| | | |
|---|---|---|
| qthdFFw | GATAATGACACTATCGTAGCCC | (4) |
| qthdFRv | GCAGTTCCAGCACATCTTC | (4) |

**Supplementary Table 2. Bioinformatic analyses supplemental data**

| Sample | B # | T # | L | Desired # of reads (M) | Obtained # of reads (M) | Trimmed reads | Reads on MG1655 (Q10) | Approximate coverage on MG1655 (x) | Reads on pVCR94 (Q10) | Coverage on pVCR94 (x) | Reads on SGI1 (Q10) | Coverage on SGI1 (x) |
|---|---|---|---|---|---|---|---|---|---|---|---|---|
| ChIPexo_sgaDC_pVCRdeltaacaDC_1 | 1 | 1 | 1 | 1 | 0,32 | 287634 | 72096 | 2 | 94018 | 114 | na | na |
| ChIPexo_sgaDC_pVCRdeltaacaDC_2 | 2 | 1 | 1 | 1 | 0,38 | 322648 | 78768 | 3 | 121940 | 147 | na | na |
| ChIPexo_sgaDC_pVCRdeltaacaDC_ SGI1deltasgaDC_1 | 1 | 1 | 1 | 1 | 0,30 | 255144 | 47288 | 2 | 68166 | 82 | 57771 | 276 |
| ChIPexo_sgaDC_pVCRdeltaacaDC_ SGI1deltasgaDC_2 | 1 | 2 | 1 | 1 | 0,34 | 308145 | 76561 | 2 | 82964 | 100 | 69895 | 334 |
| ChIPexo_sgaDC_pVCRdeltaacaDC_ SGI1deltasgaDC_3 | 2 | 1 | 1 | 1 | 0,35 | 316230 | 60961 | 2 | 92783 | 112 | 79977 | 382 |
| ChIPexo_acaDC_pVCRdeltaacaDC_ SGI1deltasgaDC_1 | 1 | 1 | 2 | 1 | 0,51 | 480195 | 48119 | 2 | 105826 | 128 | 146783 | 701 |
| ChIPexo_acaDC_pVCRdeltaacaDC_ SGI1deltasgaDC_2 | 1 | 2 | 2 | 1 | 0,51 | 478999 | 45566 | 1 | 110312 | 133 | 170019 | 812 |
| ChIPexo_acaDC_pVCRdeltaacaDC_ SGI1deltasgaDC_3 | 2 | 1 | 2 | 1 | 0,75 | 708642 | 78579 | 3 | 117709 | 142 | 196265 | 938 |
| CapSeq_sgaDC_pVCRdeltaacaDC | 1 | 1 | 1 | 1 | 1,04 | 977102 | 750797 | 24 | 19660 | 24 | na | na |
| CapSeq_pVCRdeltaacaDC | 1 | 1 | 1 | 1 | 2,39 | 2269504 | 2220355 | 72 | 5232 | 6 | na | na |
| CapSeq_acaDC_pVCRdeltaacaDC_ SGI1deltasgaDC_1 | 1 | 1 | 2 | 1 | 19,70 | 789203 | 621680 | 20 | 4978 | 6 | 6483 | 31 |
| CapSeq_acaDC_pVCRdeltaacaDC_ SGI1deltasgaDC_2 | 1 | 2 | 2 | 1 | 13,65 | 513887 | 375291 | 12 | 2420 | 3 | 7972 | 38 |
| CapSeq_pVCRdeltaacaDC_ SGI1deltasgaDC_1 | 1 | 1 | 2 | 1 | 17,62 | 849366 | 81919 | 3 | 4859 | 6 | 11052 | 53 |
| CapSeq_pVCRdeltaacaDC_ SGI1deltasgaDC_2 | 2 | 1 | 2 | 1 | 19,83 | 1052271 | 795003 | 26 | 2659 | 3 | 3152 | 15 |
| CapSeq_sgaDC_pVCRdeltaacaDC_ SGI1deltasgaDC_2 | 1 | 1 | 2 | 1 | 6,19 | 922345 | 648794 | 21 | 13476 | 16 | 13514 | 65 |
| CapSeq_sgaDC_pVCRdeltaacaDC_ SGI1deltasgaDC_3 | 2 | 1 | 2 | 1 | 7,07 | 2596089 | 1855165 | 60 | 44565 | 54 | 51553 | 246 |
| RNASeq_pVCRdeltaacaDC_2 | 1 | 1 | 1 | 5 | 9,87 | 9283646 | 5640354 | 182 | 73883 | 89 | na | na |
| RNASeq_pVCRdeltaacaDC_3 | 2 | 1 | 1 | 5 | 9,06 | 8523516 | 6158345 | 199 | 73232 | 89 | na | na |
| RNASeq_pVCRdeltaacaDC_4 | 1 | 2 | 1 | 5 | 8,89 | 8414979 | 5941814 | 192 | 102720 | 124 | na | na |

| | | | | | | | | | | | | |
|---|---|---|---|---|---|---|---|---|---|---|---|---|
| RNASeq_pVCRdeltaacaDC_5 | 2 | 2 | 1 | 5 | 9,76 | 9190911 | 6434217 | 208 | 64363 | 78 | na | na |
| RNASeq_induced_sgaDC_ pVCRdeltaacaDC_1 | 1 | 1 | 1 | 5 | 10,11 | 9597871 | 8848617 | 286 | 150922 | 182 | na | na |
| RNASeq_induced_sgaDC_ pVCRdeltaacaDC_2 | 1 | 2 | 1 | 5 | 9,42 | 8922661 | 8088735 | 261 | 181030 | 219 | na | na |
| RNASeq_induced_sgaDC_ pVCRdeltaacaDC_3 | 2 | 1 | 1 | 5 | 8,98 | 8504670 | 4800404 | 155 | 222304 | 269 | na | na |
| RNASeq_induced_sgaDC_ pVCRdeltaacaDC_4 | 2 | 2 | 1 | 5 | 7,56 | 7150962 | 4350985 | 140 | 198722 | 240 | na | na |
| RNASeq_induced_sgaDC_ pVCRdeltaacaDC_5 | 2 | 3 | 1 | 5 | 3,97 | 3741495 | 1933340 | 62 | 87977 | 106 | na | na |
| RNASeq_induced_sgaDC_ pVCRdeltaacaDC_SGI1deltasgaDC_1 | 1 | 1 | 1 | 5 | 9,33 | 8802470 | 4970555 | 160 | 277279 | 335 | 447965 | 2141 |
| RNASeq_induced_sgaDC_ pVCRdeltaacaDC_SGI1deltasgaDC_3 | 2 | 1 | 1 | 5 | 8,23 | 7778553 | 5578376 | 180 | 331062 | 400 | 469022 | 2241 |
| RNASeq_induced_sgaDC_ pVCRdeltaacaDC_SGI1deltasgaDC_4 | 2 | 2 | 1 | 5 | 8,84 | 8349834 | 5202798 | 168 | 296246 | 358 | 554795 | 2651 |
| RNASeq_induced_sgaDC_ pVCRdeltaacaDC_SGI1deltasgaDC_5 | 2 | 3 | 1 | 5 | 8,38 | 7946552 | 5473657 | 177 | 413753 | 500 | 619336 | 2960 |
| RNASeq_induced_acaDC_ pVCRdeltaacaDC_SGI1deltasgaDC_1 | 1 | 1 | 2 | 5 | 8,47 | 8198741 | 5144563 | 166 | 365399 | 442 | 309008 | 1477 |
| RNASeq_induced_acaDC_ pVCRdeltaacaDC_SGI1deltasgaDC_3 | 1 | 2 | 2 | 5 | 8,96 | 8619107 | 5226629 | 169 | 399512 | 483 | 477398 | 2281 |
| RNASeq_induced_acaDC_ pVCRdeltaacaDC_SGI1deltasgaDC_4 | 2 | 1 | 2 | 5 | 8,87 | 8594800 | 5062466 | 163 | 338885 | 410 | 481110 | 2299 |
| RNASeq_induced_acaDC_ pVCRdeltaacaDC_SGI1deltasgaDC_5 | 2 | 2 | 2 | 5 | 9,35 | 9038305 | 5045461 | 163 | 321642 | 389 | 384855 | 1839 |
| RNASeq_induced_acaDC_ pVCRdeltaacaDC_SGI1deltasgaDC_6 | 2 | 3 | 2 | 5 | 11,37 | 10958195 | 5090333 | 164 | 316293 | 382 | 367010 | 1754 |
| RNASeq_pVCRdeltaacaDC_ SGI1deltasgaDC_1 | 1 | 1 | 2 | 5 | 8,91 | 8631906 | 7449854 | 240 | 94677 | 114 | 129364 | 618 |
| RNASeq_pVCRdeltaacaDC_ SGI1deltasgaDC_2 | 1 | 2 | 2 | 5 | 8,23 | 7972683 | 7046746 | 227 | 118205 | 143 | 120607 | 576 |
| RNASeq_pVCRdeltaacaDC_ SGI1deltasgaDC_3 | 1 | 3 | 2 | 5 | 6,88 | 6682701 | 5759123 | 186 | 110485 | 134 | 196802 | 940 |
| RNASeq_pVCRdeltaacaDC_ SGI1deltasgaDC_4 | 2 | 1 | 2 | 5 | 6,92 | 6723083 | 6128823 | 198 | 65507 | 79 | 106946 | 511 |
| RNASeq_pVCRdeltaacaDC_ SGI1deltasgaDC_5 | 2 | 2 | 2 | 5 | 8,96 | 8687731 | 7982967 | 258 | 94389 | 114 | 98811 | 472 |

| | | | | | | | | | | | | |
|---|---|---|---|---|---|---|---|---|---|---|---|---|
| RNASeq_pVCRdeltaacaDC_ SGI1deltasgaDC_6 | 2 | 3 | 2 | 5 | 8,00 | 7762858 | 7093720 | 229 | 78430 | 95 | 118898 | 568 |

B; biological replicate, T; technical replicate, L; sequencing line. Samples framed in bold were chosen to represent the tracks displayed in Figure 2.

**Supplementary Table 3. Top 10 up- and down-regulated MG1655 genes upon *sgaDC* overexpression.**

| Gene | $Log_2$(fold-change) | Adjusted *p*-value | GO terms (biological process) | GO terms (molecular function) | Other notable mentions |
|---|---|---|---|---|---|
| *ydeT* | 6,9 | 2,95E-21 | GO:0055085 - transmembrane transport<br>GO:0009297 - pilus assembly | GO:0015473 - fimbrial usher porin activity | Deletion mutant shows increased biofilm formation (5) |
| *iraM* | 5,9 | 5,12E-09 | GO:0010350 - cellular response to magnesium starvation<br>GO:0071468 - cellular response to acidic pH | GO:0005515 - protein binding<br>GO:0043856 - anti-sigma factor antagonist activity | |
| *yhjB* | 5,1 | 5,91E-09 | GO:0006355 - regulation of transcription, DNA-templated | GO:0005515 - protein binding<br>GO:0003677 - DNA binding | YhjB belongs to the NarL family of response regulators (6) |
| *agaB* | 5,0 | 1,16E-05 | GO:0034219 - carbohydrate transmembrane transport | GO:0008982 - protein-N(PI)-phosphohistidine-sugar phosphotransferase activity | |
| *yaiS* | 4,9 | 2,71E-06 | | GO:0016787 - hydrolase activity | |
| *ygeF* | 4,8 | 2,33E-06 | | | Located within a remnant of an ETT2 (T3SS) pathogenicity island (7) |
| *cmtB* | 4,8 | 2,74E-06 | GO:0008643 - carbohydrate transport | GO:0016740 - transferase activity | |
| *ycgZ* | 4,7 | 3,58E-11 | GO:0071468 - cellular response to acidic pH<br>GO:0045892 - negative regulation of transcription, DNA-templated | | Overexpression increases fitness and resistance to several antibiotics (8) |
| *yihQ* | 4,7 | 1,97E-05 | GO:0005975 - carbohydrate metabolic process | GO:1990929 - sulfoquinovosidase activity<br>GO:0016798 - hydrolase activity, acting on glycosyl bonds | |

| | | | | | |
|---|---|---|---|---|---|
| *ydeO* | 4,7 | 9,69E-07 | GO:0045893 - positive regulation of transcription, DNA-templated | GO:0003700 - DNA-binding transcription factor activity | YdeO plays an important role in survival under both acidic and anaerobic conditions (9) |
| *cysI* | -5,1 | 2,60E-10 | GO:0000103 - sulfate assimilation<br>GO:0019344 - cysteine biosynthetic process | GO:0050311 - sulfite reductase (ferredoxin) activity<br>GO:0004783 - sulfite reductase (NADPH) activity | |
| *raiA* | -4,7 | 4,70E-15 | GO:0045900 - negative regulation of translational elongation<br>GO:0045947 - negative regulation of translational initiation<br>GO:0009409 - response to cold | GO:0043022 - ribosome binding | |
| *yibI* | -4,7 | 1,01E-06 | | | |
| *entD* | -4,6 | 7,82E-06 | GO:0009239 - enterobactin biosynthetic process | GO:0008897 - holo-[acyl-carrier-protein] synthase activity | |
| *garD* | -4,5 | 2,95E-21 | GO:0046392 - galactarate catabolic process | GO:0008867 - galactarate dehydratase activity<br>GO:0016829 - lyase activity | |
| *nmpC* | -4,5 | 4,57E-07 | | | Encodes a porin associated with increased sensitivity to several colicins and phages (10) |
| *narGHJI* | -4,3 | 7,31E-16 | GO:0009061 - anaerobic respiration<br>GO:0042128 - nitrate assimilation | GO:0009055 - electron transfer activity<br>GO:0008940 - nitrate reductase activity | |
| *nrfA* | -4,3 | 1,39E-19 | GO:0019645 - anaerobic electron transport chain<br>GO:0042128 - nitrate assimilation | GO:0042279 - nitrite reductase (cytochrome, ammonia-forming) activity | |

| | | | | |
|---|---|---|---|---|
| *nirD* | -4,3 | 3,29E-05 | GO:0009061 - anaerobic respiration<br>GO:0042128 - nitrate assimilation | GO:0008942 - nitrite reductase [NAD(P)H] activity |
| *cysA* | -4,2 | 4,19E-05 | GO:0015709 - thiosulfate transport<br>GO:1902358 - sulfate transmembrane transport | GO:0102025 - ATPase-coupled thiosulfate transmembrane transporter activity |

**Supplementary Table 4. Predicted AcaCD/SgaCD-responsive promoters in other families of conjugative plasmids.**

| Plasmid | Accession # | Gene/Locus tag[a] | Start | Stop | Strand | Score[b] | *p*-value[b] | *q*-value[b] | Matched sequence[b] |
|---|---|---|---|---|---|---|---|---|---|
| pAsa4c | NZ_KT033470.1 | - | 12326 | 12353 | - | 27,0102 | 8,64E-10 | 4,69E-05 | AAACTGCCCGGATTGGGCACCAACACCG |
| pAsa4c | NZ_KT033470.1 | *traI* | 36563 | 36590 | + | 31,5000 | 3,28E-11 | 5,34E-06 | AAACTGCCCAAATTGGACAGTTACGCAG |
| pAsa4c | NZ_KT033470.1 | - | 43729 | 43756 | - | 25,3061 | 2,38E-09 | 9,06E-05 | TTTGCGCCCAATTCGGGCAGTTACAGCG |
| pAsa4c | NZ_KT033470.1 | *traL* | 43734 | 43761 | + | 23,0816 | 7,83E-09 | 2,55E-04 | TAACTGCCCGAATTGGGCGCAAAGCCGT |
| pAsa4c | NZ_KT033470.1 | *traV* | 46946 | 46973 | + | 30,5612 | 7,11E-11 | 7,72E-06 | ATTGTGACCAAAATGGGCAGTTTCACAG |
| pAsa4c | NZ_KT033470.1 | *dsbC* | 53669 | 53696 | + | 32,3980 | 1,48E-11 | 4,81E-06 | GATGTGACCAAAATGGGCCGTTACACAG |
| pAsa4c | NZ_KT033470.1 | HTG22_RS00315 | 66272 | 66299 | + | 21,9286 | 1,39E-08 | 4,11E-04 | AAGTTGCCCAAAATGGACGTCTACAGCG |
| pAsa4c | NZ_KT033470.1 | HTG22_RS00370 | 76581 | 76608 | + | 25,6122 | 2,00E-09 | 9,06E-05 | ATTTCACCCGAAAAGGGCAGCTTGAGCG |
| pAsa4c | NZ_KT033470.1 | HTG22_RS00460 | 89711 | 89738 | + | 27,1735 | 7,80E-10 | 4,69E-05 | TTTGCGCCCCAAAAAGGCAGTTACAGCG |
| pAsa4c | NZ_KT033470.1 | *traF* | 150655 | 150682 | + | 25,2143 | 2,50E-09 | 9,06E-05 | AATGCGCCCTTAATGGGCGCCAACACCT |
| pAsa4c | NZ_KT033470.1 | *parM* | 161774 | 161801 | - | 27,8878 | 4,92E-10 | 4,01E-05 | TAAATGCCCATTTTGGGCAGTTCCAGAG |
| pVPS91 | NZ_KX957972.1 | *traV* | 10285 | 10312 | + | 22,816 | 8,95E-09 | 5,83E-04 | AAGCTGACCAGAAAGGGCAGTTTCACAG |
| pVPS91 | NZ_KX957972.1 | HTL46_RS00070 | 11772 | 11799 | + | 29,622 | 1,46E-10 | 2,38E-05 | AAAACGCTCAAATTGGGCAGTTACACAG |
| pVPS91 | NZ_KX957972.1 | *dsbC* | 17409 | 17436 | + | 22 | 1,34E-08 | 7,27E-04 | TAATTGACCAAACAGGGCAACTACAGAT |
| pVPS91 | NZ_KX957972.1 | *traN* | 24534 | 24561 | + | 20,878 | 2,28E-08 | 9,29E-04 | GTGTTGTCCAAAATGGACAGTTCCACAG |
| pVPS91 | NZ_KX957972.1 | HTL46_RS00310 | 58179 | 58206 | + | 20,939 | 2,22E-08 | 9,29E-04 | CAAACGCCCTAAAAGGGCAAATAGACGT |
| pVPS91 | NZ_KX957972.1 | HTL46_RS00450 | 79965 | 79992 | + | 24,959 | 2,89E-09 | 2,35E-04 | AAACTGCCCAAAATGGACCATTTCACGA |
| pVPS91 | NZ_KX957972.1 | HTL46_RS00450 | 80013 | 80040 | + | 18,99 | 5,34E-08 | 1,74E-03 | TATTTGCCCAAAATGGGCTGATGCAGAG |
| pVPS91 | NZ_KX957972.1 | *traF* | 83903 | 83930 | + | 20,286 | 3,00E-08 | 1,08E-03 | TAATTGCCCTAAATGGGAGGTTTCAGAT |
| pVPS91 | NZ_KX957972.1 | HTL46_RS00640 | 113346 | 113373 | + | 25,449 | 2,19E-09 | 2,35E-04 | TATGCGCCCGTATAGGGCAGTTAAGCAT |
| pVPS91 | NZ_KX957972.1 | HTL46_RS00920 | 156001 | 156028 | + | 34,959 | 1,02E-12 | 3,33E-07 | ATTGCGCCCAAAAAGGGCAGTTACAGGT |
| pVPS91 | NZ_KX957972.1 | *traI* | 162964 | 162991 | + | 18,5 | 6,61E-08 | 1,96E-03 | AAACTGCCCAAAATGGACAGGAGGGGAG |
| pAhD4-1 | NZ_CP013966.1 | AhyD4_RS23175 | 38016 | 38043 | - | 24,776 | 3,20E-09 | 1,25E-04 | TTTGCACCCCAAAAAGGCAGTTACAGCG |
| pAhD4-1 | NZ_CP013966.1 | AhyD4_RS23255 | 50883 | 50910 | - | 27,99 | 4,60E-10 | 2,87E-05 | ATTGCACCCGAAAAGGGCAGTTTGAGCG |
| pAhD4-1 | NZ_CP013966.1 | AhyD4_RS23310 | 61021 | 61048 | - | 24,225 | 4,31E-09 | 1,49E-04 | AAGTTGCCCAAAATGGACTGTTACAGCG |

| | | | | | | | | | |
|---|---|---|---|---|---|---|---|---|---|
| pAhD4-1 | NZ_CP013966.1 | *dsbC* | 73058 | 73085 | - | 30,449 | 7,77E-11 | 1,10E-05 | GATGTGACCAAATTGGGCCGTTACAGAG |
| pAhD4-1 | NZ_CP013966.1 | *traV* | 79773 | 79800 | - | 33,755 | 3,90E-12 | 1,21E-06 | ATTGTGACCAAAAAGGGCAGTTACACAG |
| pAhD4-1 | NZ_CP013966.1 | *traL* | 82976 | 83003 | - | 23,806 | 5,38E-09 | 1,68E-04 | TAACTGCCCTAAATGGGCGCAAAGCCGT |
| pAhD4-1 | NZ_CP013966.1 | - | 82981 | 83008 | + | 29,541 | 1,55E-10 | 1,21E-05 | TTTGCGCCCATTTAGGGCAGTTACAGCG |
| pAhD4-1 | NZ_CP013966.1 | *traI* | 93232 | 93259 | - | 30,051 | 1,06E-10 | 1,10E-05 | AAACTGCCCAAATTGGACAGTTAGGCAG |
| pAhD4-1 | NZ_CP013966.1 | - | 120376 | 120403 | + | 25,959 | 1,63E-09 | 8,48E-05 | GAACTGCCCGGATTGGGCACCAACAGAG |
| pAhD4-1 | NZ_CP013966.1 | *traF* | 145826 | 145853 | - | 25,214 | 2,50E-09 | 1,11E-04 | AATGCGCCCTTAATGGGCGCCAACACCT |
| pAhD4-1 | NZ_CP013966.1 | - | 145831 | 145858 | + | 19,122 | 5,04E-08 | 1,43E-03 | TTGGCGCCCATTAAGGGCGCATTGAGCG |
| pAQU1 | NC_016983.1 | PAQU1_002 | 1486 | 1513 | + | 25,449 | 2,19E-09 | 2,23E-04 | TATGCGCCCGTATAGGGCAGTTAAGCAT |
| pAQU1 | NC_016983.1 | *traI* | 48352 | 48379 | + | 18,5 | 6,61E-08 | 2,69E-03 | AAACTGCCCAAAATGGACAGGAGGGGAG |
| pAQU1 | NC_016983.1 | *traV* | 58679 | 58706 | + | 22,816 | 8,95E-09 | 7,30E-04 | AAGCTGACCAGAAAGGGCAGTTTCACAG |
| pAQU1 | NC_016983.1 | PAQU1_074 | 60167 | 60194 | + | 29,622 | 1,46E-10 | 5,97E-05 | AAAACGCTCAAATTGGGCAGTTACACAG |
| pAQU1 | NC_016983.1 | *dsbC* | 65753 | 65780 | + | 22 | 1,34E-08 | 9,10E-04 | TAATTGACCAAACAGGGCAACTACAGAT |
| pAQU1 | NC_016983.1 | *traN* | 72878 | 72905 | + | 20,878 | 2,28E-08 | 1,16E-03 | GTGTTGTCCAAAATGGACAGTTCCACAG |
| pAQU1 | NC_016983.1 | PAQU1_128 | 113823 | 113850 | + | 20,939 | 2,22E-08 | 1,16E-03 | CAAACGCCCTAAAAGGGCAAATAGACGT |
| pAQU1 | NC_016983.1 | - | 171835 | 171862 | + | 28,51 | 3,24E-10 | 6,60E-05 | AAACTGCCCAAAATGGACCATTACACGA |
| pAQU1 | NC_016983.1 | PAQU1_198 | 171883 | 171910 | + | 27,765 | 5,33E-10 | 7,25E-05 | TATTTGCCCAAAATGGGCTGATACAGAG |
| pAQU1 | NC_016983.1 | *traF* | 175774 | 175801 | + | 20,286 | 3,00E-08 | 1,36E-03 | TAATTGCCCTAAATGGGAGGTTTCAGAT |

[a] Gene or locus tag upstream of which the predicted AcaCD box was detected. "-" indicates that no gene or locus tag was found immediately upstream on the same strand.

[b] Predictions of AcaCD-binding site were carried out by using FIMO (Find Individual Motif Occurrences) V5.1.1 (11) with the AcaCD MEME logo, as described elsewhere (2). Description of the score, *q*- and *p*-values is available at http://alternate.meme-suite.org//doc/fimo-output-format.html.

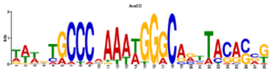

```
vcrx012 ctgcag(41)-GGGTTGCCCGAATTGGGCAGCTAGGCGGGGTTGCATTTTTGAAAAGTCCTGTACGCTGGTTGGTATA(14)-ctcgagcaggaaacagctATG
vcrx035 ctgcag(95)-TACGCGCCCATTTTGGGCAGATAGACAAGTTATTTGGGCGTTTGGAGTGTGCGAGGTGGACAGCGGG(156)ctgcagattcgactcgagcaggaaacagctATG
vcrx036 ctgcag(200)TATCTGCCCAAAATGGGCGCGTAAGCGTCCGATAGAAAGTAATTGTTAGTGAGATATTGACTACACG(51)-ctgcagattcgactcgagcaggaaacagctATG
vcrx059 ctgcag(57)-TATCTGTCCAATTTGGGCAGTTTGAGAAACCAGCTTTCCACCCCCCCCCTTTTAGAATGGGCTCATC(14)-ctcgagcaggaaacagctATG
traI ctgcag(58)-AAACTGCCCAAATTGGACAGATAGGCGTGTGTGCGATCTAAAGGTGTTTTGAGAGCATGTAGCGACG(13)-ctgcagattcgactcgagcaggaaacagctATG
vcrx068 ctgcag(7)--AAAGTGCCCAAAATGGGCCGTAACAGCACTGTTCCTAGATTGATCAATTAAGAGACGATTACTCCGT(27)-ctcgagcaggaaacagctATG
traL ctgcag(56)-GAAGTGCCCGAAAAGGGCGCGAAGCCGTGGCAGAAAATCAGTGACCCTGAATGAGAATACAGGCCAC(11)-ctgcagattcgactcgagcaggaaacagctATG
traA ctgcag(208)ATTTTGACCAGAAAGGGCAGTTCCAGAGGCGAGGCTCAAAAGGAACAGGCAGTAAGATCCGCAAAGA(36)-ctcgagcaggaaacagctATG
vcrx076 ctgcag(99)-TATTTGCTCAAAAAGGGCACTTCCACAGCCCATGTTTATGCACTCCCTCCCGTAACCTCCAGCATTG(33)-ctcgagcaggaaacagctATG
dsbC ctgcag(54)-GAATTGTCCAAATTGGGCTATTACAGAGGCGAGTCTATTTCCCTCCCCCGGATACACTGCCCAGCAT(27)-ctcgagcaggaaacagctATG
traN_C ctgcag(33)-ATTATGCCCCAAAAGGGCCACTACACGTACAGGTGGCCCTTGTCTTCTCAGATAGCATGGGAAGGTT(15)-ctcgagcaggaaacagctATG
acaB ctgcag(192)TAACTGTCCATTTTGGGCACTTCGAGATGTCAGGTTACAGATTGACCGCATTTATCCTTTCCAGTGC(33)-ctgcagattcgactcgagcaggaaacagctATG
vcrx087 ctgcag(77)-GAAGTGCCCAAAATGGACAGTTACAGCGTTTTCTGTTGAGCAAAGGTGAGCTATATTTAAGCCATC(149)-ctgcagattcgactcgagcaggaaacagctATG
vcrx098 ctgcag(13)-TTTGCACCCGAAAAGGGCAGTTTGAGCGTTCCTGTTCACACAGCTCTCGTGGAACAATCAAAGTGAG(184)ctcgagcaggaaacagctATG
vcrx114 ctgcag(28)-TTTGCACCCCAAAAGGGCAATTACAGCGTTTTACCCTTTTAGGTGGCTGTCATACTCTTTGCTCAT(107)-ctcgagcaggaaacagctATG
vcrx128 ctgcag(50)-TTCATACCCTAAAAGGGCAAAAACGCAGTCGTTGATAACGACTCTCCCACCTGAAAATAGCTCCAG(185)-ctcgagcaggaaacagctATG
vcrx140 ctgcag(38)-TTTGAGCCAAAAATTGGCAACTACAGAGGGTGAGTTGGAGGAGCAAAAAGGCAAGAGTATAGAGCAG(54)-ctcgagcaggaaacagctATG
traF ctgcag(46)-TAAGCGCCCTAAATGGGCGCTTACACCTGCCTAATTTCACGCCTCCCACCTCTACCATATCGAGCATGG---ctgcagattcgactcgagcaggaaacagctATG
vcrx152 ctgcag(368)TATATGCCCCTTTTGGGCAATTACACAGGCCATCTGTCCCCGCTTTTAGCCTTATTCTCTATCCAAC(41)-ctgcagattcgactcgagcaggaaacagctATG
xis ctgcag(253)TTCGCGCCCTAAAAGGGCAGATCCAGAGCCGAGGATTGGTAGCTGTCTCTCATAAACTAGGTCTATCA(5)-ctgcagattcgactcgagcaggaaacagctATG
S004 ctgcag(120)TAGGTGCCCAAATAGGGCACTTCCAGAGGCCATGTTTACGGGGGTTCACCCTCAAAATTTCTCCAGT(28)-ctcgagcaggaaacagctATG
traN_S ctgcag(109)CGTATGCCGCAAAAGGGCAAATAGCGATGCTAATTTTTATGAGAGAGCGACATAGCATTATCCAAAC(11)-ctgcagattcgactcgagcaggaaacagctATG
traH_S ctgcag(152)GGACTGCCCAAATTGGACAGTTTGGGAGTTCGGTTTGTACTCATTTGCTCGGTAGGCTTTCGGGTGA(11)-ctgcagattcgactcgagcaggaaacagctATG
S018 ctgcag(74)-AATGTGCCCAAAAAGGGCCAATACAGCGCGTGTCGATTCAGAAATGAGCTTGTTAAATTTGAATCA(123)-ctgcagattcgactcgagcaggaaacagctATG
```

**Supplementary Figure 1. Content of pOP*lac*Z-derivative vectors used for β-galactosidase assays.** For each vector, the cloned insert corresponds to the sequence between the two restriction sites (either PstI or XhoI) written in orange. AcaCD binding site is written in green. TSS is written in blue and was updated from Carraro *et al*, 2014 (2), using the Cappable-seq data obtained in this study. Predicted -10 and -35 motifs are indicated in purple and red, respectively. Promoters that displayed a constitutive activity are indicated in brown.

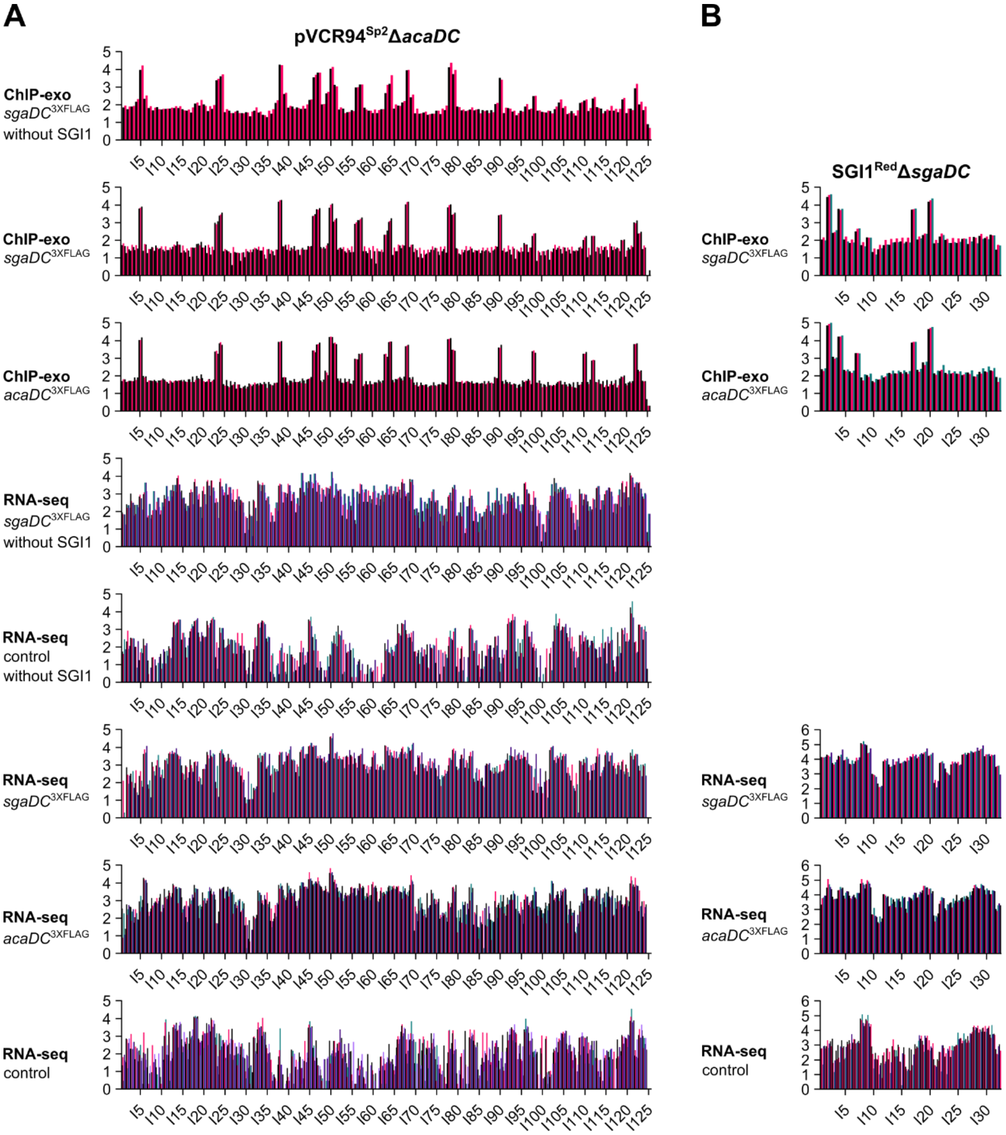
A
pVCR94Sp2ΔacaDC
ChIP-exo
sgaDC3XFLAG
without SGI1
ChIP-exo
sgaDC3XFLAG
ChIP-exo
acaDC3XFLAG
RNA-seq
sgaDC3XFLAG
without SGI1
RNA-seq
control
without SGI1
RNA-seq
sgaDC3XFLAG
RNA-seq
acaDC3XFLAG
RNA-seq
control
B
SGI1RedΔsgaDC
ChIP-exo
sgaDC3XFLAG
ChIP-exo
acaDC3XFLAG
RNA-seq
sgaDC3XFLAG
RNA-seq
acaDC3XFLAG
RNA-seq
control

C

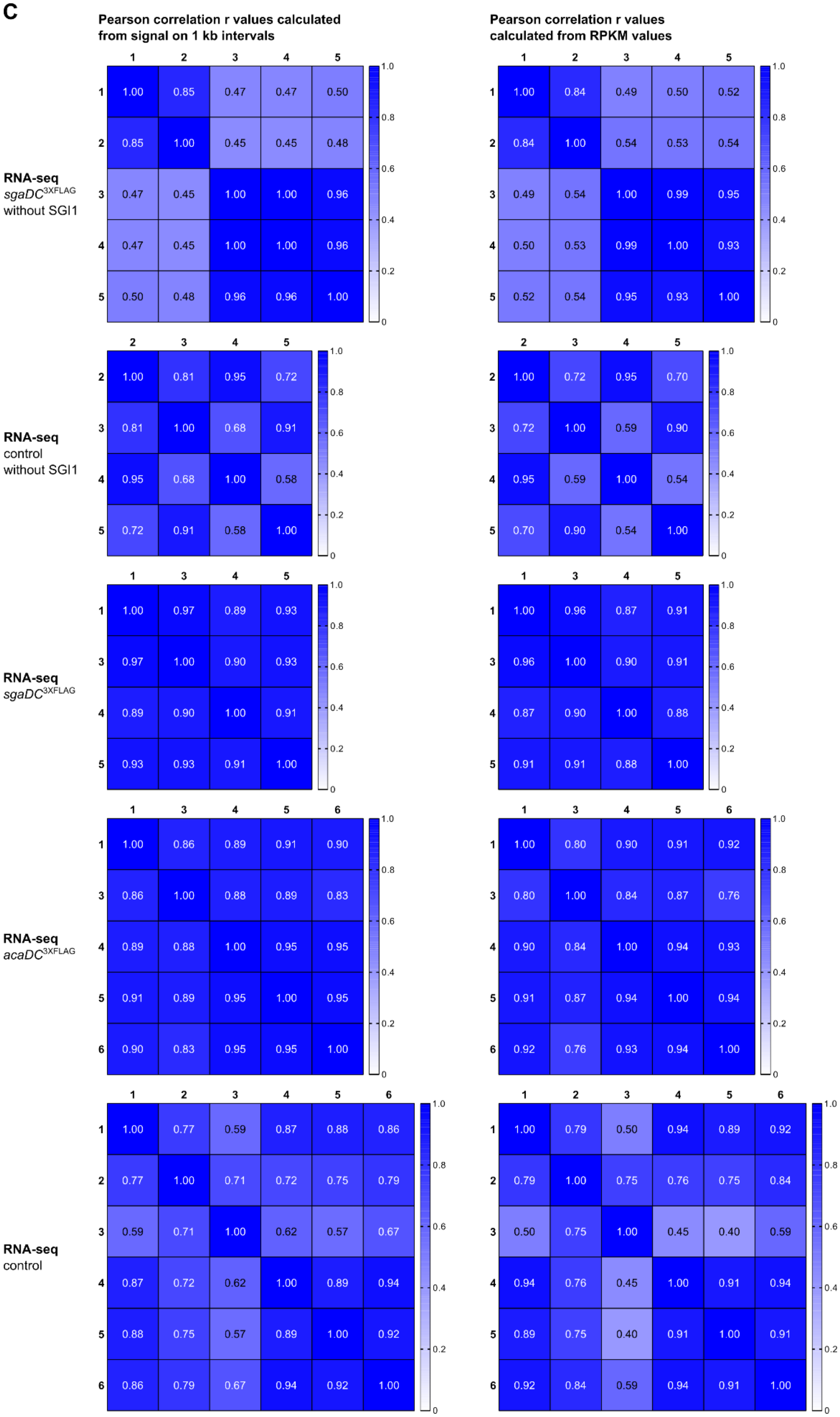
Pearson correlation r values calculated from signal on 1 kb intervals
Pearson correlation r values calculated from RPKM values
RNA-seq sgaDC3XFLAG without SGI1
1 2 3 4 5
1 1.00 0.85 0.47 0.47 0.50
2 0.85 1.00 0.45 0.45 0.48
3 0.47 0.45 1.00 1.00 0.96
4 0.47 0.45 1.00 1.00 0.96
5 0.50 0.48 0.96 0.96 1.00
1 2 3 4 5
1 1.00 0.84 0.49 0.50 0.52
2 0.84 1.00 0.54 0.53 0.54
3 0.49 0.54 1.00 0.99 0.95
4 0.50 0.53 0.99 1.00 0.93
5 0.52 0.54 0.95 0.93 1.00
RNA-seq control without SGI1
2 3 4 5
2 1.00 0.81 0.95 0.72
3 0.81 1.00 0.68 0.91
4 0.95 0.68 1.00 0.58
5 0.72 0.91 0.58 1.00
2 3 4 5
2 1.00 0.72 0.95 0.70
3 0.72 1.00 0.59 0.90
4 0.95 0.59 1.00 0.54
5 0.70 0.90 0.54 1.00
RNA-seq sgaDC3XFLAG
1 3 4 5
1 1.00 0.97 0.89 0.93
3 0.97 1.00 0.90 0.93
4 0.89 0.90 1.00 0.91
5 0.93 0.93 0.91 1.00
1 3 4 5
1 1.00 0.96 0.87 0.91
3 0.96 1.00 0.90 0.91
4 0.87 0.90 1.00 0.88
5 0.91 0.91 0.88 1.00
RNA-seq acaDC3XFLAG
1 3 4 5 6
1 1.00 0.86 0.89 0.91 0.90
3 0.86 1.00 0.88 0.89 0.83
4 0.89 0.88 1.00 0.95 0.95
5 0.91 0.89 0.95 1.00 0.95
6 0.90 0.83 0.95 0.95 1.00
1 3 4 5 6
1 1.00 0.80 0.90 0.91 0.92
3 0.80 1.00 0.84 0.87 0.76
4 0.90 0.84 1.00 0.94 0.93
5 0.91 0.87 0.94 1.00 0.94
6 0.92 0.76 0.93 0.94 1.00
RNA-seq control
1 2 3 4 5 6
1 1.00 0.77 0.59 0.87 0.88 0.86
2 0.77 1.00 0.71 0.72 0.75 0.79
3 0.59 0.71 1.00 0.62 0.57 0.67
4 0.87 0.72 0.62 1.00 0.89 0.94
5 0.88 0.75 0.57 0.89 1.00 0.92
6 0.86 0.79 0.67 0.94 0.92 1.00
1 2 3 4 5 6
1 1.00 0.79 0.50 0.94 0.89 0.92
2 0.79 1.00 0.75 0.76 0.75 0.84
3 0.50 0.75 1.00 0.45 0.40 0.59
4 0.94 0.76 0.45 1.00 0.91 0.94
5 0.89 0.75 0.40 0.91 1.00 0.91
6 0.92 0.84 0.59 0.94 0.91 1.00
1.0
0.8
0.6
0.4
0.2
0

**Supplementary Figure 2. Correlation between ChIP-exo and RNA-seq replicates.** ChIP-exo and RNA-seq signals for each replicate (both strands) are represented as a log-transformed function of the position in (A) pVCR94$^{Sp2}$ Δ*acaDC* or (B) SGI1$^{Red}$ Δ*sgaDC*, where both elements have been split in 1-kb intervals (I1, I2, etc). For additional details, refer to legend of Fig 2. (C) RNA-seq signal obtained with pVCR94$^{Sp2}$ Δ*acaDC* was used to compare the replicates for each indicated condition (detailed in Table S2). Pearson correlation r values calculated either from signal on 1 kb intervals (panels on the left) or RPKM values (panels on the right) are shown as heatmaps.

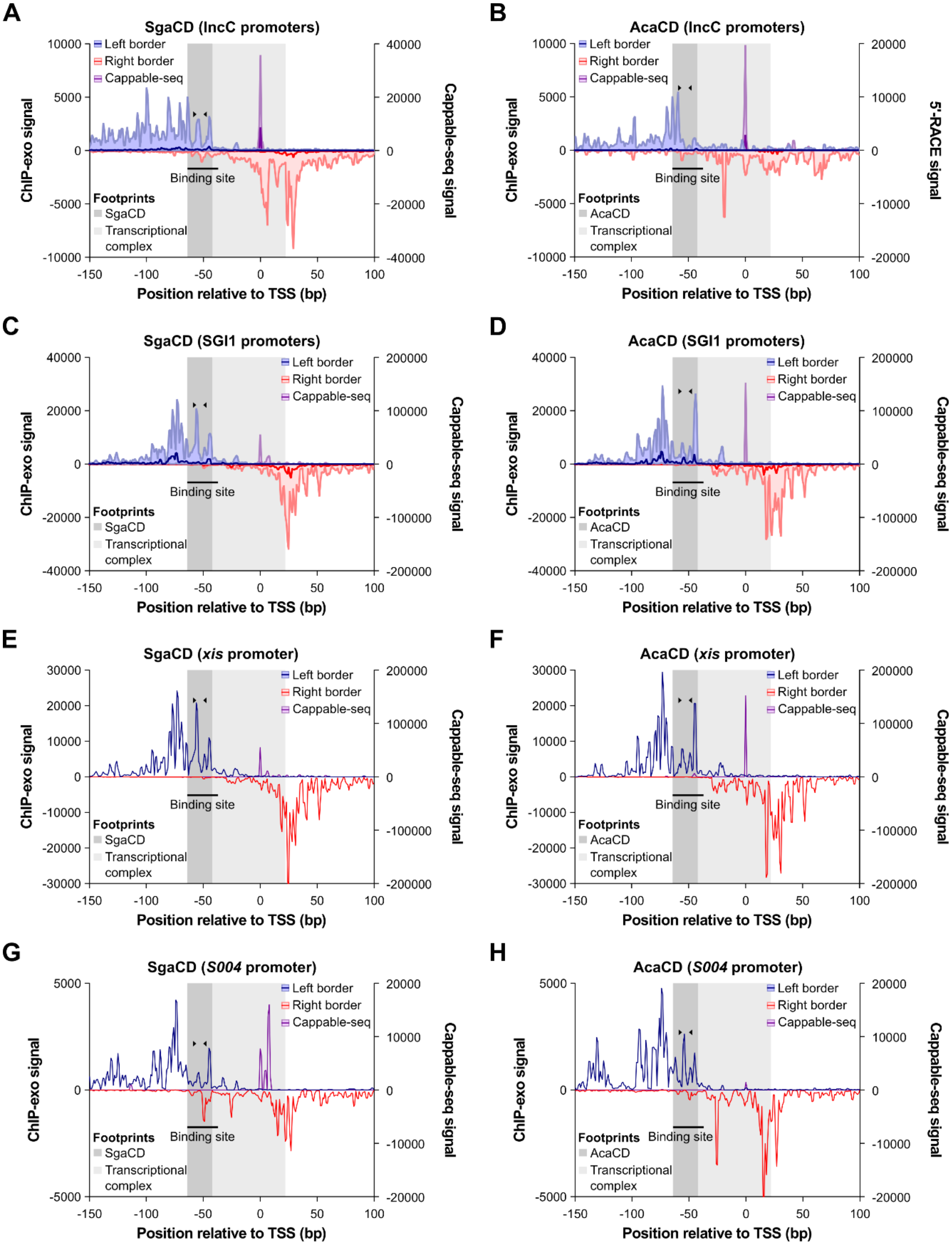

**Supplementary Figure 3. VAP aggregate profiles of AcaCD/SgaCD footprint.** ChIP-exo and Cappable-seq profiles obtained with SgaCD (A) and ChIP-exo and 5'-RACE profiles obtained with AcaCD (B) were aggregated from 13 AcaCD-activatable IncC promoters. ChIP-exo and Cappable-seq profiles obtained with SgaCD (C) and AcaCD (D) were aggregated from 5 SGI1 AcaCD-activatable promoters. Individual profiles are shown for *xis* (E-F) and *S004* promoters (G-H) The normalized density of ChIP-exo reads mapping on the positive DNA strand (left border, in blue) and the negative strand (right border, in red), as well as the normalized density of Cappable-seq or 5'-RACE reads mapping on the positive strand (purple), are plotted as functions of the distance in nucleotides to the aggregated transcription start site (TSS). Data are represented as median and range. The binding site is indicated as a black line, with its **GCCC**NDWWW**GGGC** motif delimitated by black triangles. The region protected by the transcriptional complex, delimitated by the last and first peaks on the ChIP-exo left and right borders, respectively, is depicted in light grey. The region protected by either SgaCD or AcaCD, starting at the binding site, is depicted in dark grey.

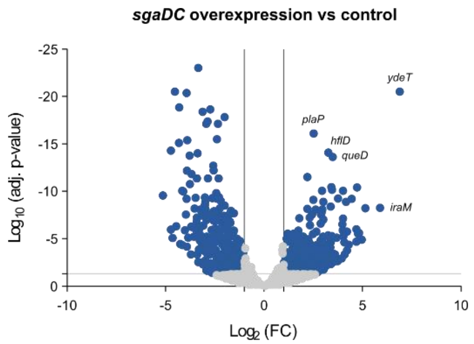

**Supplementary Figure 4. Differential expression analysis.** Volcano plot of *E. coli* MG1655 genes using the $Log_2$ of fold-change and adjusted *p*-value. The vertical lines indicate a 2-fold change in expression used as the threshold, and the horizontal line defines statistical significance at $p = 0.05$. Differentially expressed genes are marked as blue dots.

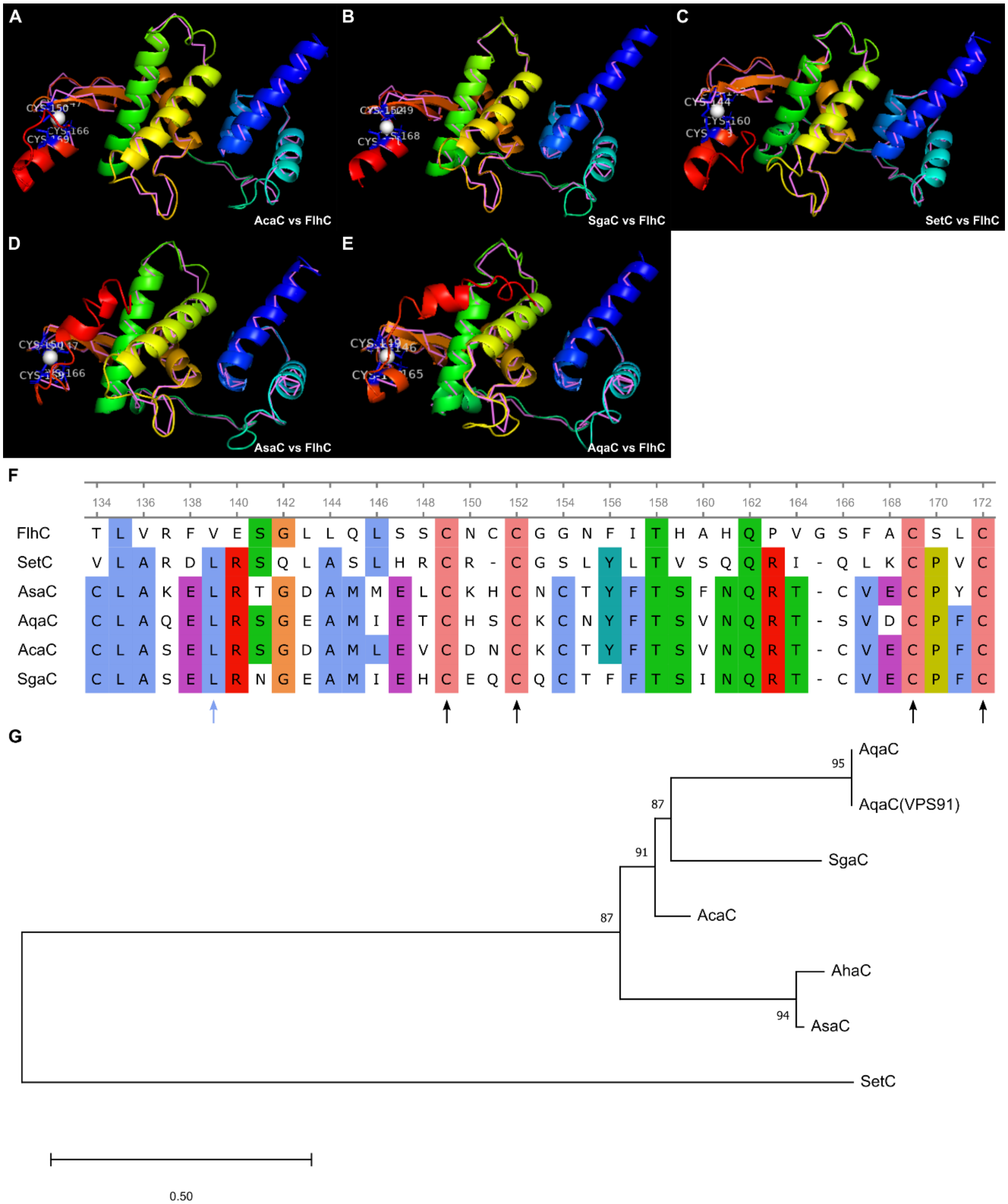

**Supplementary Figure 5. Predicted structures of AcaC, SgaC, SetC, AsaC and AqaC.** (A-E) Predicted structures of AcaC, SgaC, SetC, AsaC and AqaC. Query and FlhC structures are shown in cartoon and pink backbone, respectively. Structures were predicted by I-TASSER (12). PDB 2AVU:E (FlhC) was specified as template (without alignment) and was identified as having the closest structural similarity (i.e. the highest TM-score) to the query predicted model. $Zn^{2+}$ (white sphere) was identified as the most probable ligand by COFACTOR and COACH (13, 14). Corresponding binding cysteines are displayed in blue sticks and labeled. (F) Alignment of the primary sequence of the zinc-finger domain of the AcaC orthologues, displaying an almost conserved leucine residue at position 139, as well as the four conserved cysteines involved in the predicted zinc-finger domain. (G) Phylogenetic analysis of AcaC orthologues inferred by using the Maximum Likelihood method and Whelan And Goldman model (15). Primary sequences were aligned with MUSCLE (16). The tree with the highest log likelihood is shown. Initial tree(s) for the heuristic search were obtained automatically by applying Neighbor-Join and BioNJ algorithms to a matrix of pairwise distances estimated using the JTT model, and then selecting the topology with superior log likelihood value. A discrete Gamma distribution was used to model evolutionary rate differences among sites (5 categories (+*G*, parameter = 0,9821)). The tree is drawn to scale, with branch lengths measured in the number of substitutions per site. There was a total of 201 positions in the final dataset. Evolutionary analyses were conducted in MEGA X (17). AsaC (ALL42451.1), AqaC (WP_014386839.1), AqaC (VPS91) (WP_014386839.1) and AhaC (WP_047235007.1) are encoded by pAsa4c from *Aeromonas salmonicida* subsp. *salmonicida* (18), pAQU1 from *Photobacterium damselae* subsp. *damselae* (19), pVPS91 from *Vibrio parahaemolyticus* (20), and pAhD4-1 from *Aeromonas hydrophila* (21), respectively.

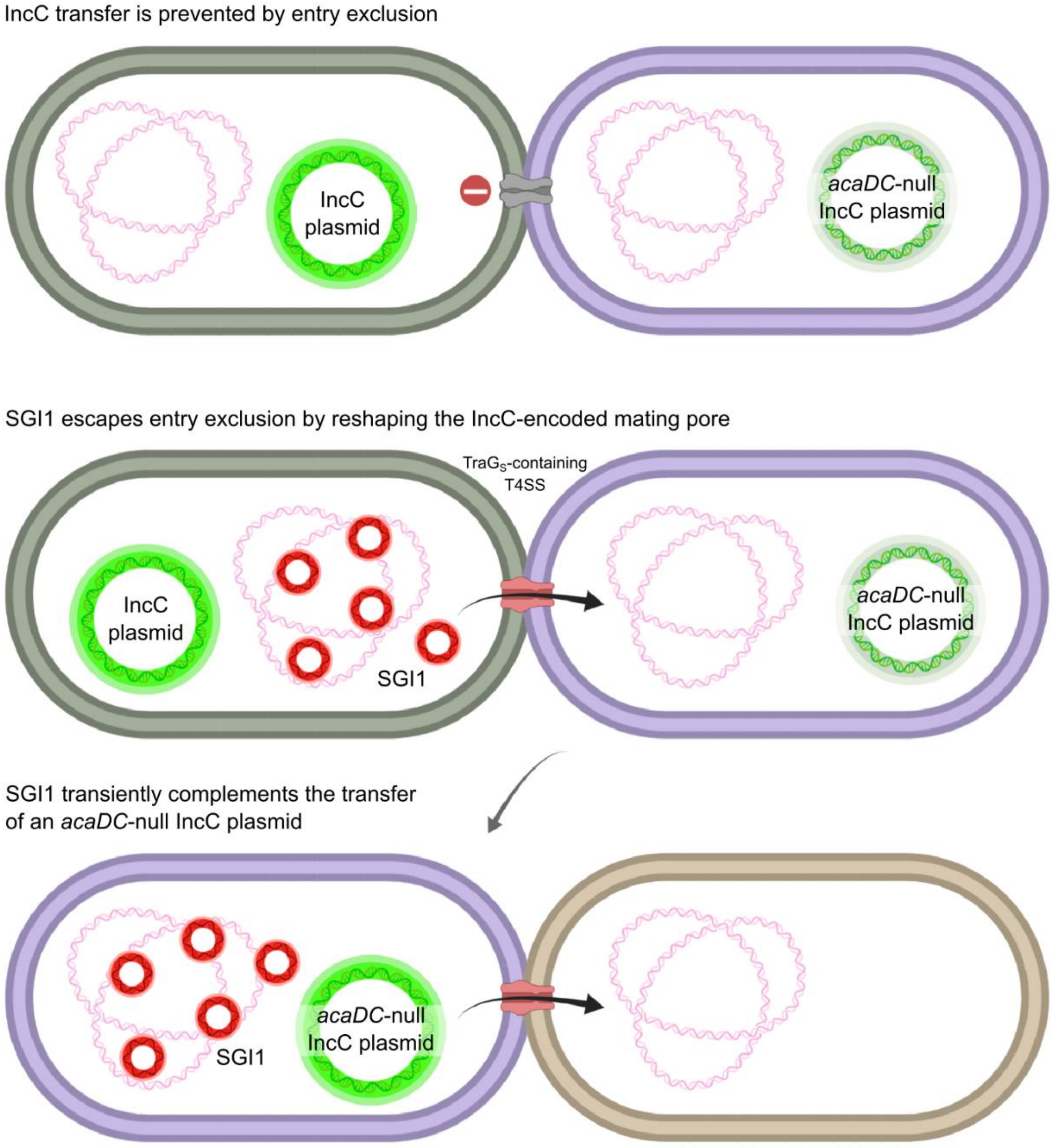

**Supplementary Figure 6. Interactions between SGI1 and IncC plasmids.** Drawings illustrate the incapacity of an IncC plasmid to transfer to a cell already bearing an IncC plasmid, because of entry exclusion (top) and the escape by SGI1 of such mechanism, allowed by the replacement of TraG$_C$ by TraG$_S$ (middle). Following SGI1 transfer to a strain already containing an *acaDC*-null IncC plasmid, SGI1 is able to complement the transfer of said plasmid to another strain (bottom). Created with BioRender.com.

### References

1. Carraro,N., Sauvé,M., Matteau,D., Lauzon,G., Rodrigue,S. and Burrus,V. (2014) Development of pVCR94ΔX from *Vibrio cholerae*, a prototype for studying multidrug resistant IncA/C conjugative plasmids. *Front Microbiol*, **5**.

2. Carraro,N., Matteau,D., Luo,P., Rodrigue,S. and Burrus,V. (2014) The Master Activator of IncA/C Conjugative Plasmids Stimulates Genomic Islands and Multidrug Resistance Dissemination. *PLoS Genet*, **10**, e1004714.

3. Carraro,N., Durand,R., Rivard,N., Anquetil,C., Barrette,C., Humbert,M. and Burrus,V. (2017) *Salmonella* genomic island 1 (SGI1) reshapes the mating apparatus of IncC conjugative plasmids to promote self-propagation. *PLOS Genetics*, **13**, e1006705.

4. Huguet,K.T., Rivard,N., Garneau,D., Palanee,J. and Burrus,V. (2020) Replication of the *Salmonella* Genomic Island 1 (SGI1) triggered by helper IncC conjugative plasmids promotes incompatibility and plasmid loss. *PLOS Genetics*, **16**, e1008965.

5. Vlasblom,J., Zuberi,K., Rodriguez,H., Arnold,R., Gagarinova,A., Deineko,V., Kumar,A., Leung,E., Rizzolo,K., Samanfar,B., *et al.* (2015) Novel function discovery with GeneMANIA: a new integrated resource for gene function prediction in *Escherichia coli*. *Bioinformatics*, **31**, 306–310.

6. Yamamoto,K., Hirao,K., Oshima,T., Aiba,H., Utsumi,R. and Ishihama,A. (2005) Functional Characterization in Vitro of All Two-component Signal Transduction Systems from *Escherichia coli*. *J. Biol. Chem.*, **280**, 1448–1456.

7. Ren,C.-P., Chaudhuri,R.R., Fivian,A., Bailey,C.M., Antonio,M., Barnes,W.M. and Pallen,M.J. (2004) The ETT2 gene cluster, encoding a second type III secretion system from *Escherichia coli*, is present in the majority of strains but has undergone widespread mutational attrition. *J. Bacteriol.*, **186**, 3547–3560.

8. Soo,V.W.C., Hanson-Manful,P. and Patrick,W.M. (2011) Artificial gene amplification reveals an abundance of promiscuous resistance determinants in *Escherichia coli*. *Proc. Natl. Acad. Sci. U.S.A.*, **108**, 1484–1489.

9. Yamanaka,Y., Oshima,T., Ishihama,A. and Yamamoto,K. (2014) Characterization of the YdeO regulon in *Escherichia coli*. *PLoS ONE*, **9**, e111962.

10. Pugsley,A.P. and Schnaitman,C.A. (1978) Identification of three genes controlling production of new outer membrane pore proteins in *Escherichia coli* K-12. *J. Bacteriol.*, **135**, 1118–1129.

11. Grant,C.E., Bailey,T.L. and Noble,W.S. (2011) FIMO: scanning for occurrences of a given motif. *Bioinformatics*, **27**, 1017–1018.

12. Yang,J., Yan,R., Roy,A., Xu,D., Poisson,J. and Zhang,Y. (2015) The I-TASSER Suite: protein structure and function prediction. *Nature Methods*, **12**, 7–8.

13. Zhang,C., Freddolino,P.L. and Zhang,Y. (2017) COFACTOR: improved protein function prediction by combining structure, sequence and protein–protein interaction information. *Nucleic Acids Res*, **45**, W291–W299.

14. Yang,J., Roy,A. and Zhang,Y. (2013) Protein–ligand binding site recognition using complementary binding-specific substructure comparison and sequence profile alignment. *Bioinformatics*, **29**, 2588–2595.

15. Whelan,S. and Goldman,N. (2001) A General Empirical Model of Protein Evolution Derived from Multiple Protein Families Using a Maximum-Likelihood Approach. *Mol Biol Evol*, **18**, 691–699.

16. Edgar,R.C. (2004) MUSCLE: multiple sequence alignment with high accuracy and high throughput. *Nucleic Acids Res*, **32**, 1792–1797.

17. Kumar,S., Stecher,G., Li,M., Knyaz,C. and Tamura,K. (2018) MEGA X: Molecular Evolutionary Genetics Analysis across Computing Platforms. *Mol Biol Evol*, **35**, 1547–1549.

18. Tanaka,K.H., Vincent,A.T., Trudel,M.V., Paquet,V.E., Frenette,M. and Charette,S.J. (2016) The mosaic architecture of *Aeromonas salmonicida* subsp. *salmonicida* pAsa4 plasmid and its consequences on antibiotic resistance. *PeerJ*, **4**, e2595.

19. Nonaka,L., Maruyama,F., Miyamoto,M., Miyakoshi,M., Kurokawa,K. and Masuda,M. (2012) Novel Conjugative Transferable Multiple Drug Resistance Plasmid pAQU1 from *Photobacterium damselae* subsp. *damselae* Isolated from Marine Aquaculture Environment. *Microbes and Environments*, **27**, 263–272.

20. Humbert,M., Huguet,K.T., Coulombe,F. and Burrus,V. (2019) Entry Exclusion of Conjugative Plasmids of the IncA, IncC, and Related Untyped Incompatibility Groups. *J Bacteriol*, **201**.

21. Zhu,L., Zheng,J.-S., Wang,W.-M. and Luo,Y. (2019) Complete Genome Sequence of Highly Virulent *Aeromonas hydrophila* Strain D4, Isolated from a Diseased Blunt-Snout Bream in China. *Microbiol Resour Announc*, **8**.
